# Axonal Branch Points of Parvalbumin Interneurons are Focal Sites of Tau-Induced Degeneration

**DOI:** 10.64898/2026.09.17.751394

**Authors:** Paul Feyen, Yuxiao Zhang, Theresa Niedermeier, Nicolas Landgraf, Florence M Bareyre, Carsten Vos, Lars Paeger, Jochen Herms

## Abstract

Tau aggregation, a defining feature of tauopathies, commonly appears as neuropil threads within neurites including axons. However, the affected neuronal populations and the spatial organization of axonal vulnerability remain poorly defined. Here, following identification of Tau accumulation in parvalbumin (PV) interneuron axons in primary tauopathy patients, we generated a P301S Tau model targeted to PV interneurons to track the structural integrity of their neurites and somas. We observed neuronal loss alongside axonal dystrophy, with focal swellings forming preferentially at axonal bifurcations. This spatial enrichment exceeded chance levels, and local axonal geometry predicted swelling localization. Longitudinal in vivo imaging revealed that bifurcation-associated swellings progressed to axonal severing. Analogous bifurcation-associated dystrophies were present in PV axons in human primary tauopathy tissue. These findings indicate that axonal branch geometry shapes the spatial pattern of Tau-induced neurodegeneration and identify PV axonal branch points as sites of pathology in mouse and human tauopathy.

## Introduction

Tauopathies are a group of neurodegenerative diseases characterized by pathological aggregation of the microtubule-associated protein Tau in distinct cellular subtypes and cellular sub-compartments (*1*, *2*). A frequent feature across tauopathies is the presence of ‘neuropil threads’, accumulations of Tau proteins in the neurites of neurons. Extensive Tau-positive neuritic networks and bundles of affected axons are seen in subcortical gray and white-matter tracts in progressive supranuclear palsy (PSP) (*3*), and neuropil-thread pathology is abundant in cortical gray and white matter in corticobasal degeneration (CBD) (*1*, *4*). In the secondary tauopathy Alzheimer’s disease (AD), pretangle Tau pathology and Tau-positive neuritic pathology emerge early in the transentorhinal cortex and become widespread in the entorhinal region before overt neocortical neurofibrillary tangle (NFT) pathology is observed (*5*, *6*). Despite ample evidence of axonal Tau accumulation across diverse tauopathies, the molecular identity of the afflicted neurons remains largely undefined and the consequences of axonal Tau accumulation on axonal viability are poorly understood. Additionally, axons comprise multiple sub-compartments, including synapses, varicosities and branch points, any of which could be differentially susceptible to Tau-induced damage by virtue of their geometry and molecular composition.

A prerequisite of defining neuronal subtypes bearing neuropil threads is the ability to identify molecular identity and simultaneously assess for Tau inclusions. In this work, we focus on parvalbumin (PV) expressing interneurons of the cortex, whose neurites can be visualized by the broad cytosolic distribution of the PV protein. PV neurons are known for their extensively branched and myelinated axonal arbor, fast firing rates, and central roles in network oscillations (*7*, *8*). These features impose strong demands on cytoskeletal organization, axonal transport, and metabolic support, all of which are processes with documented disturbance in tauopathy models (*9–11*). Moreover, evidence from single-cell sequencing of AD tissue sorted by AT8 levels (an antibody recognizing phosphorylated Tau) revealed that 6.8% of PV+ chandelier neurons harbor NFTs in the neocortex (*12*), with further histological identification of PV+ neuron Tau aggregation in the hippocampal subgranular zone and hilus (*13*). Despite these observations, it remains unclear whether PV neuron axons themselves develop Tau proteinopathy, whether such axonal lesions contribute directly to structural or functional deficits, and whether PV Tau aggregation extends beyond secondary tauopathy (AD) to primary tauopathies such as CBD and PSP. We therefore aimed to assess the presence and eventual consequences of PV axo-dendritic Tau load in the brain of primary tauopathy patients and in a cell-type restricted mouse model.

Following identification of neuritic PV tauopathy in CBD patients, we generated a PV-restricted 2N4R P301S Tau expression model using AAVs in PV-Cre mice. Here PV neurons expressed full length human Tau bearing a mutation identified in a family with history of FTD and CBD (*14*). We observed pronounced PV neuron loss and focal axonal dystrophy that was enriched at axon bifurcation points with distinct geometric signatures, suggesting a microdomain-specific vulnerability within PV neurons. Longitudinal *in vivo* two-photon microscopy revealed accompanying oligodendrocyte reactivity and bifurcation-centered swellings that progressed to axon severing. Analogous PV⁺ bifurcation-associated dystrophies were identified in post-mortem brain tissue of human primary tauopathy patients. Collectively, these findings indicate that axonal damage in tauopathy is spatially related to axonal branch geometry, and that PV axonal branch points represent vulnerable loci in mouse and human tauopathy.

## Results

### Parvalbumin neuron soma and axons present with Tau accumulation in primary tauopathy

To assess for Tau proteinopathy in PV somas and neurites, we performed immunohisto-fluorescence (IHF) in free floating tissue sections prepared from the Superior Frontal Gyrus of CBD Donors (Fig. 1). The cell type, Tau accumulation, and axonal compartment, were identified with antibodies against PV, pTau (AT8), and Myelin Basic Protein (MBP) respectively. We set out to answer three questions; Do PV somas bear tau inclusions? Do PV neurites bear tau inclusions, and if so are those neurites axons? In all tested cases, PV somas bearing characteristic AT8+ Tau threads and pre-threads were identified (Fig1.b-d, N = 4, Sup.Table1). The locations of AT8-positive somas broadly followed the global distribution of PV somas across the cortex, indicating that somatic Tau inclusions can affect PV neurons across a large range of cortical depths (Fig.1c,d). At a population level, ∼1.9-3.3% of PV neuron somas had Tau inclusions.

**Figure 1.**
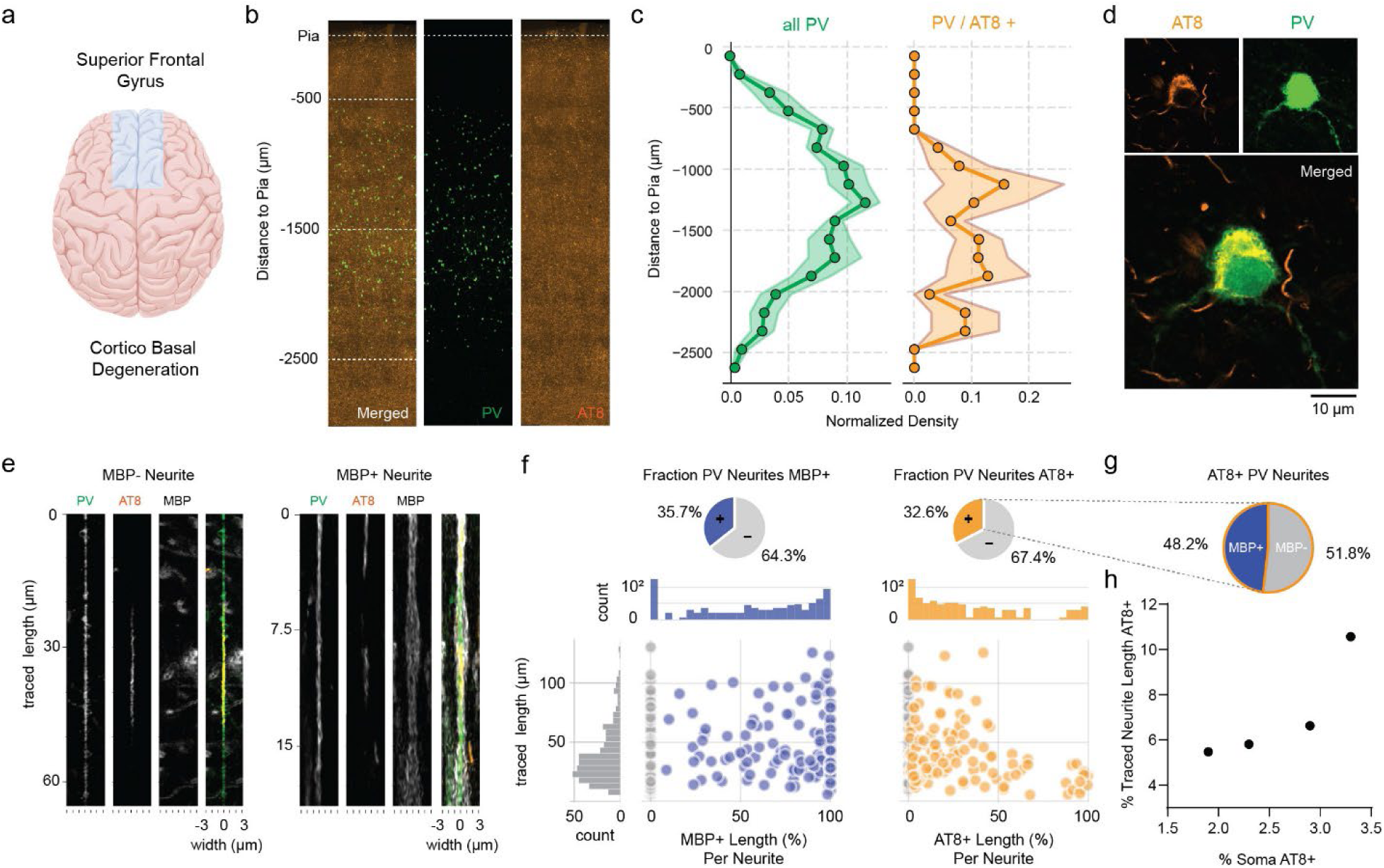
Parvalbumin neurons present with Tau accumulation in primary tauopathy. **(a)** 100 µm sections from the superior frontal gyrus of CBD patients were stained via IHF for Parvalbumin, AT8 and MBP. **(b)** Example image showing distribution of PV and AT8 signal in cortex from Pia to white matter. **(c)** Normalized density (mean ± SEM) as a function of distance from pia of PV somas (green) and PV somas bearing Tau inclusions denoted by AT8 (orange) **(d)** High magnification image of an AT8 positive parvalbumin soma. **(e)** Examples of digitally straightened Parvalbumin neurites bearing neuropil threads in a non-myelinated neurite segment (left) and a myelinated neurite (right). **(f)** Fraction of neurites that were MBP positive (blue) and fraction of neurites that were AT8 positive (orange). Lower panels show corresponding quantification of the analyzed PV neurites, showing lengths of analyzed neurite segments in relation to the percent of the neurite’s length that is myelinated or AT8 positive. **(g)** Fraction of AT8-positive neurites that were MBP-positive or MBP-negative **(h)** Scatter plot of the % of traced neurite length that is AT8 positive and the percent of AT8 positive soma per analyzed case.

To determine if PV neurites presented with proteinopathy, we acquired 3D super-resolution Airyscan stacks of triple labeled sections (example single plane, Sup.Fig.1). This permitted the analysis of individual neurites (Fig.1e-h). Neurites were classified as axons if they colocalized with MBP signal. Neurites without MBP coverage in contrast reflect either unmyelinated axonal segments or dendritic processes. In total, 16,049 µm of PV neurites were traced, from a mean ± SEM of 105 ± 4.8 neurites per case (Fig. 1f). Across cases, 35.7% of traced neurites were identified as axons, corresponding to 33.9% of the total traced neurite length being myelinated. Similarly, 32.6% of traced neurites were AT8-positive, corresponding to 7.6% of the total traced neurite length (Fig. 1f). Among all AT8-positive neurites, 48.2% were also MBP-positive (Fig. 1g), indicating that tau-positive PV neurites include myelinated axonal segments. This AT8/MBP double-positive population was observed in all cases (Sup.Fig.2), and yielded a confirmed axonal identity between 19 and 72% of AT8-positive PV neurites across the assessed cases. In sum, we find that PV neuron somas and neurites, including myelinated axonal segments, develop Tau proteinopathy in human primary tauopathy. To our knowledge, this represents the first direct attribution of neuropil threads to neurites of a molecularly defined neuronal subtype in human primary tauopathy.

### Parvalbumin tauopathy model develops phosphorylated Tau, neuritic swellings, and cell loss

Tauopathies occur in a dense setting of multiple cellular players. To delineate elements that are cell-autonomous and downstream of the observed PV tauopathy, we utilized AAVs in combination with a Cre-recombinase driver line to generate a cell-type restricted PV-tauopathy model. Mice were injected either with a MAPT-P301S construct in which the red fluorescent protein mKate2 was linked via a P2A sequence, or with a control construct expressing mKate2 alone (Fig.2a). Injections were targeted to the secondary motor cortex (Fig2.b,c), an area with high transcriptional correspondence to the human superior frontal gyrus (*15*). In both groups, mean specificity and local transduction efficiency exceeded 90%, as assessed by somatic colocalization of Parvalbumin protein and mKate2 (Fig.2d,e). Expression of the MAPT-P301S construct gave rise to hyper-phosphorylated and oligomeric Tau as early as one-month post-injection, an effect absent in the control group (Fig.2f,g). By six months post-injection, MAPT-P301S mice showed extensive loss of PV neurons when assessed via PV or mKate2 immunolabeling (Sup.Fig.3).

**Figure 2.**
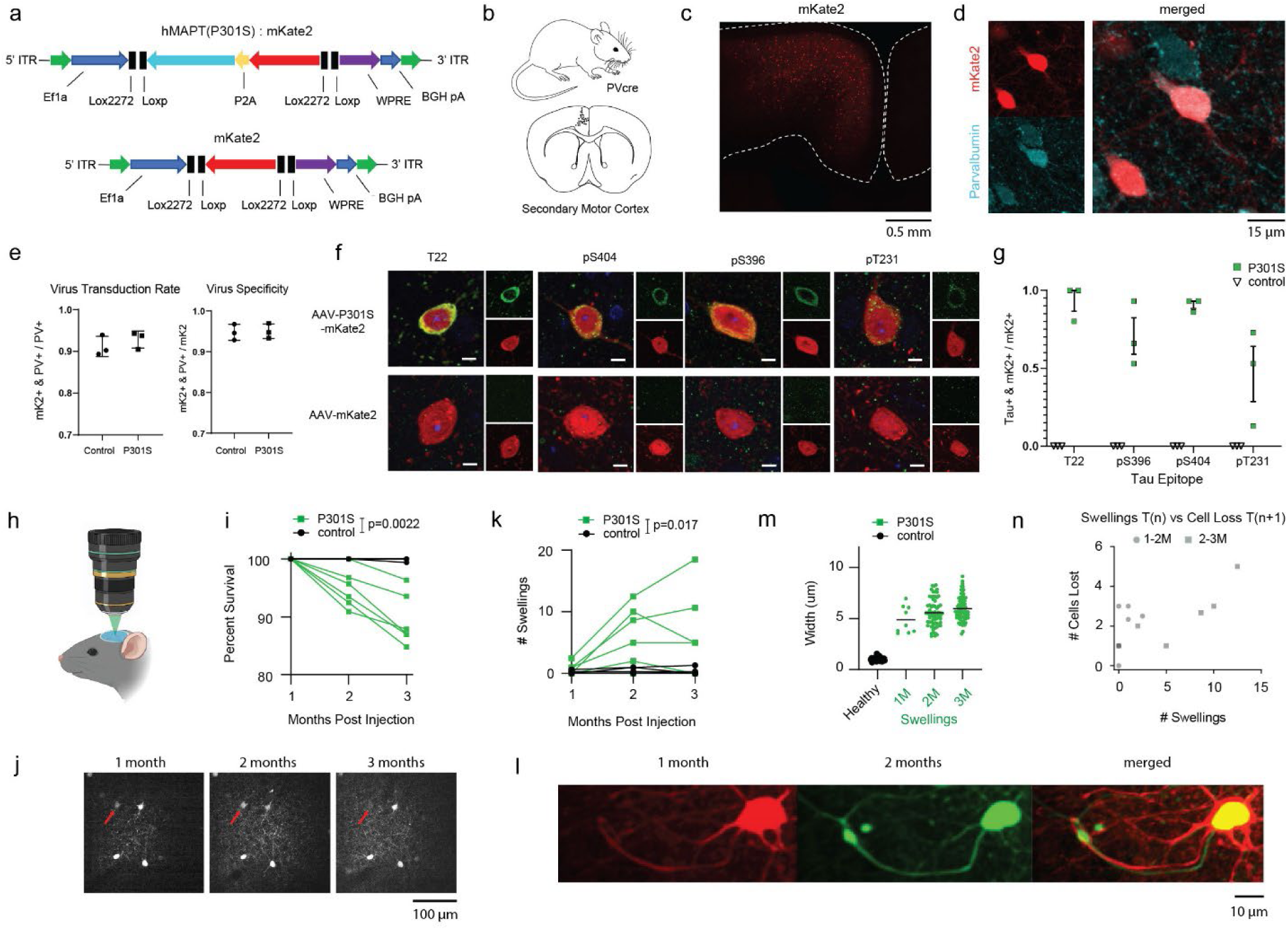
Parvalbumin-specific tauopathy model develops phosphorylated Tau and focal neuritic swellings associated with cell loss. **(a)** Virally delivered constructs to induce Tauopathy or serve as controls were injected in **(b)** the secondary motor cortex of PV-Cre mice. **(c)** Overview image with cortical boundary denoted by white dashed line showing the delivery and expression of constructs. **(d)** Close up image showing expression of mKate2 within parvalbumin neurons. **(e)** Quantification of viral transduction rate and virus specificity (N=3 mice per group). **(f)** Example images of PV neurons 1 month after expression showing presence of oligomeric Tau (T22) and hyperphosphorylated Tau protein (pS404, pS396, pT231) in mice injected with Tauopathy construct but not in controls (scale bars = 5µm). Green channel showing Tau markers, red channel showing mKate2 confirming transduction of neuron. **(g)** Per subject fraction of transduced neurons (mKate2 positive) positive for respective Tau markers (N=3 mice per group). **(h)** Longitudinal 2-photon imaging via cranial window was carried out to monitor neuronal viability and neuritic alterations. **(i)** Quantification of population survival per mouse across timepoints, control N = 6, P301S N = 6. For each mouse, longitudinal change was summarized as the mean survival at months 2 and 3 relative to the month 1 baseline, which was defined as 100%. Changes from baseline were compared between groups using a two-sided exact permutation Mann-Whitney test, p = 0.0022. **(j)** Example of a single optical slice from a longitudinally imaged volume, red arrow pointing to PV neuron loss at 3^rd^ timepoint. **(k)** Quantification of neuritic swelling burden per subject across timepoints, control N = 6, P301S N = 6. For each mouse, longitudinal change was summarized as the mean swelling burden at months 2 and 3 relative to its month 1 baseline. Changes from baseline were compared between groups using a two-sided exact permutation Mann-Whitney test, p = 0.017. **(l)** Example of a Neuritic swelling which developed between the first (red, 1-month post injection) and second (green, 2-months post injection) imaging timepoint. **(m)** Quantification of healthy neurite diameter and the diameter of observed swellings, black bar indicating mean (control: n = 50 neurites; P301S: n_total_ = 156 swellings). **(n)** Scatter plot with number of swellings at timepoint N versus PV soma loss between N and N+1. Each point represents the mean for one subject-interval pair for the 1 to 2 and the 2 to 3 month intervals from P301S group. Associations were assessed separately for each interval using one-sided Spearman rank correlations: 1M-2M, ρ = 0.360, p = 0.24 & 2M-3M, ρ = 0.899, p = 0.0074.

We next employed longitudinal in vivo two-photon imaging to assess whether Tau accumulation was associated with neuritic pathology and neuronal loss. The same cortical volumes were imaged longitudinally over three months, and individual PV somas were tracked across sessions. Compared with controls, PV neurons expressing the Tau transgene showed progressively reduced viability over the imaging period (Fig. 2h-j). Longitudinal change from baseline differed significantly between groups (Fig. 2i), resulting in a mean survival at three months of 99.9% in controls versus 89.4% in P301S mice.

Tau-expressing PV neurons also developed prominent neuritic swellings (Fig. 2k,l), defined as focal enlargements exceeding threefold the normal neurite diameter. Swellings were already detectable at the first imaging time point and increased markedly over time in the P301S group, with longitudinal change from baseline differing significantly between groups (Fig. 2k). The mean diameters of the swellings were 4.87 ± 0.44 µm, 5.56 ± 0.16 µm, and 5.97 ± 0.13 µm at one, two, and three months post-injection respectively, compared with 1.05 ± 0.02 µm diameters of healthy neurites in controls (Fig.2m). We next asked whether neuritic dystrophy burden preceded subsequent PV soma loss. Based on the hypothesis that greater swelling burden would be associated with greater cell loss at the following imaging session, we tested this relationship separately for the two longitudinal intervals (Fig. 2n). The association was weak during the 1M-2M interval (Spearman ρ = 0.360, p = 0.242) but became pronounced during the 2M-3M interval (ρ = 0.899, p = 0.0074). Thus, the association between dystrophy burden and subsequent neuronal loss emerged most clearly at the later stage of disease progression.

### Neuritic swellings in the PV-P301S-Tau mice are axonal, accumulate organelles and have a strong association to axonal bifurcations

Having found that Tau accumulation in PV neurons leads to progressive neuritic dystrophy and cell loss, we sought to define the structural identity and subcellular features of these swellings. In *ex vivo* slice preparations, dystrophic neurites were visualized via mKate2 expression, and could at times be traced back to their originating and morphologically intact parent somas, providing a direct view of these lesions within the neurite arbor of individual PV neurons (Fig. 3a). To determine whether the swellings correspond to axonal or dendritic dystrophy, we co-stained for myelin basic protein (MBP) and mKate2 in *ex vivo* brain slices of mice expressing MAPT-P301S for one month. The large majority of swellings occurred in MBP positive neurites (55/58 swellings, 94.8%; 90.9-100% across mice; N = 3), indicating pronounced axonal dystrophy in the PV tauopathy model (Fig.3b-d). We further found that a large proportion of the swellings included axonal bifurcations, denoted by the fact that 47/58 swellings (81.0%; 75.0-86.4% across mice) had three or more discernible emanating branches (Fig.3d). Swellings along a single branch segment of an axon would have only two associated branches, a single entry and exit branch. A 3D reconstruction of a bifurcation associated axonal swelling, traceable to its originating soma is presented in Supplementary Figure 4. Measuring the maximum width of mature myelinated dystrophies from three and six months post injection subjects revealed that myelin diameter was locally expanded, in tight correlation with the axonal swelling diameter (Fig.3e,f, and example 3D reconstructed bifurcation centered swelling in Sup. Video 1). We next checked whether the swellings represent sites of local organelle accumulation, a common feature in neuritic dystrophy across various etiologies. We found that lysosomes, mitochondria, and also proteins as denoted by mKate2 accumulated locally at the swelling sites (Fig.3g,h). Kif5A and parvalbumin protein showed similar accumulation in the dystrophies (Fig.3i,j), together suggesting a local deregulation of intracellular transport and/or diffusion.

**Figure 3.**
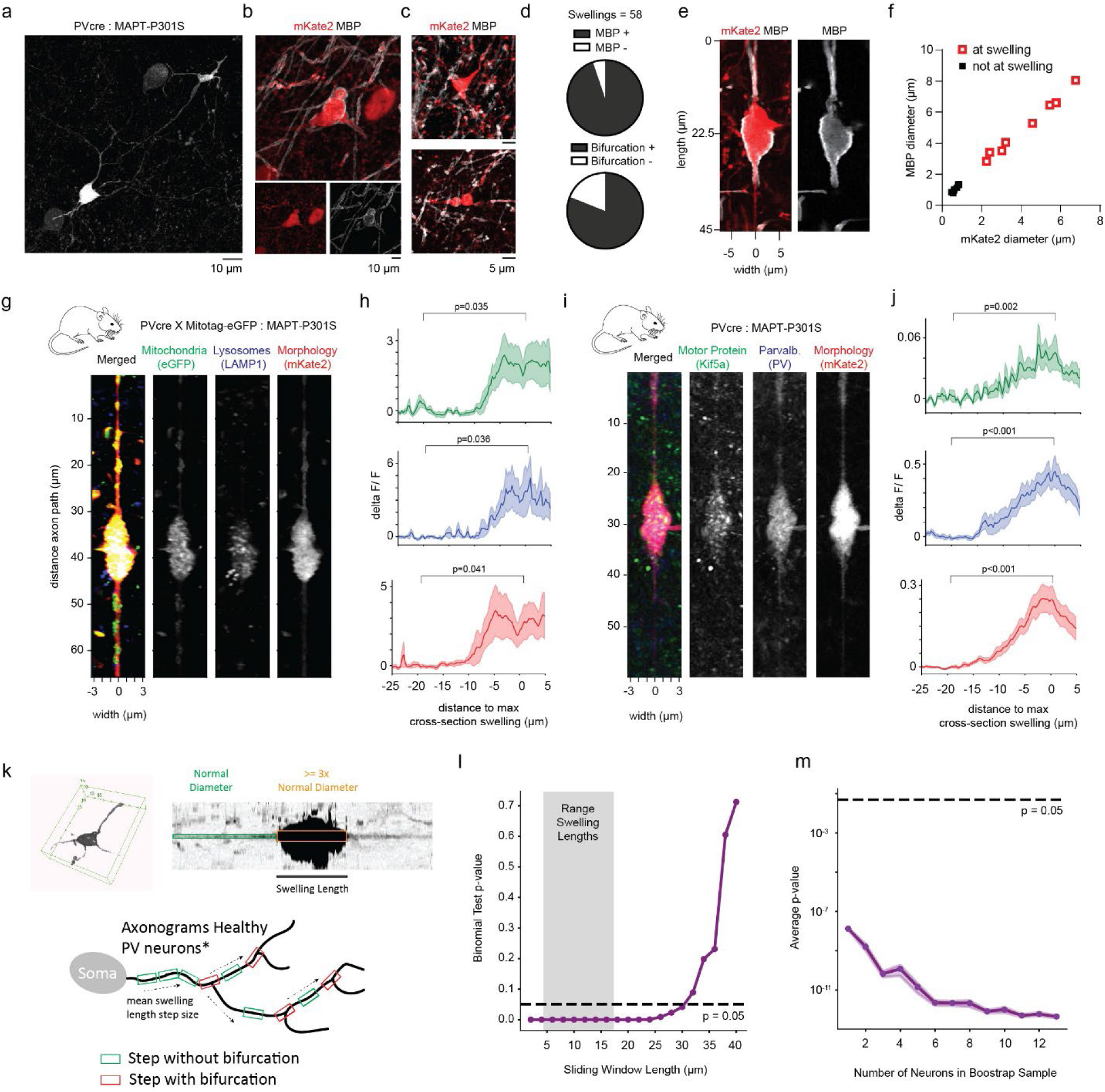
Neuritic swellings in the PV-MAPT-P301S mice are axonal, accumulate organelles and have a strong association to axonal bifurcations. **(a)** Example PV neuron expressing MAPT-P301S bearing a dystrophy (top right of image) along a neurite projecting from the soma (bottom left). Neuron imaged via mKate2 fluorophore reporter. **(b)** Example dystrophy identifiable as an axon via MBP coverage. **(c)** Two further example images of dystrophies with multiple emanating branches and MBP positive neuritic segments. **(d)** Fraction of swellings occurring along MBP positive or negative neurite, and the fraction of swellings with >=3 branches emanating (N=3 mice). **(e)** Example of a straightened axon swelling and **(f)** Quantification of the relationship between axon diameter and MBP diameter for positions inside and outside of swellings (red and black markers respectively, inside: r^2^=0.989, p<0.0001; outside: r^2^=0.931, p=0.0001). **(g-j)** Organelle and protein accumulation at PV neurite swellings. PV-Cre × MitoTag-eGFP mice were used to visualize mitochondria and were immunostained for LAMP1 to label lysosomes; separate PV-Cre mice were immunostained for Kif5a and PV (n = 7 swellings, N = 2 mice and n = 11 swellings, N = 3 mice, respectively). Line plots show fluorescence intensity measured within a 250-nm-wide region centered on the axon path, reflecting accumulation of mitochondria, lysosomes, Kif5a, parvalbumin, and mKate2 (mean ± SEM). Fluorescence intensity increased significantly from baseline (−25 to −15 µm) to the swelling region (−5 to +5 µm) for mitochondria-eGFP, lysosomes-LAMP1, Parvalbumin, Kif5a, and mKate2 (see respective p-values in figure). **(k)** (upper) Example 3D rendering of an imaged PV axonal swelling, and a straightened view of the same axon along the longest path through the swelling used to determine ‘swelling length’. (lower) Schematic of the modeling approach used to determine whether the association of swellings with branching points is a chance event based on the measured PV swelling lengths, and known cortical PV axonal arbor geometries from publicly available datasets (n=13 axon arbors, *axonograms from Micheva et al, 2021). **(l)** P-value of binomial test as a function of sliding window length utilized, with grey highlighted region showing the measured range of swelling lengths from n=10 swellings. **(m)** Average p-value of binomial test as a function of included PV neuron axonograms.

The observation that most neuritic swellings were situated at axonal bifurcations raised the possibility that branching points represent structurally vulnerable sites within the PV neuron axonal arbor. To test whether this association could arise by chance, we compiled published reconstructions of cortical parvalbumin-axon arbors (*16*) and linearized each axonal path from soma to axon terminal at 0.1-µm resolution, marking branch points. To estimate the probability that a randomly positioned axonal segment would encompass a bifurcation, we moved a virtual window along each axon path, with window lengths matched to empirically measured swelling lengths (Fig.3k, mean swelling length 11.69 µm; range 4.5-17.2 µm). This probability was p ≈ 0.34 across PV neuron axon reconstructions. We compared this baseline to our *ex vivo* data, where 47 of 58 swellings occurred at bifurcations (∼81%; 95% CI = 0.69-0.89) and a one-sided binomial test indicated that the observed proportion was far higher than expected by chance (p < 0.001 for mean swelling length). The enrichment remained robust under bootstrapping across PV-axonograms and across simulated swelling lengths: sweeping window sizes from 2-40 µm yielded strong significance throughout the observed range of swelling lengths (Fig.3l,m). These analyses demonstrate that the clustering of dystrophies at bifurcations is not a chance event, and reveal axonal branching points as focal sites of structural vulnerability in the PV-P301S tauopathy model.

### Dystrophies of axonal bifurcations can occur at different branch order points along the axonal arbor and their locations can be predicted by local axon arbor geometry

We next aimed to determine further structural characteristics of the bifurcation dystrophies. To this end, we used patch-clamp mediated biocytin filling to generate 3D reconstructions of PV axon arbors bearing dystrophies (Fig. 4a). Following cell filling, slices were fixed and stained with streptavidin-alexa conjugates to visualize morphology and immunostained against MBP to identify axonal branches. PV neurons were filled at one and three months post injection. The approach also provided a secondary somatic readout, with PV-P301S neurons showing depolarized resting membrane potentials at three months (Sup. Fig.5). From a total of 37 patched P301S neurons, ten presented with extensive axonal arbor in the slice and a focal neuritic swelling, and could be used for morphological reconstruction. All the swellings were located in the axonal arbor, as confirmed by MBP positivity (n = 3 at one month, n = 7 at three months post injection). An example biocytin filled PV neuron, along with an inset of the bifurcation associated axonal dystrophy is presented in Figure 4a-b.

**Figure 4.**
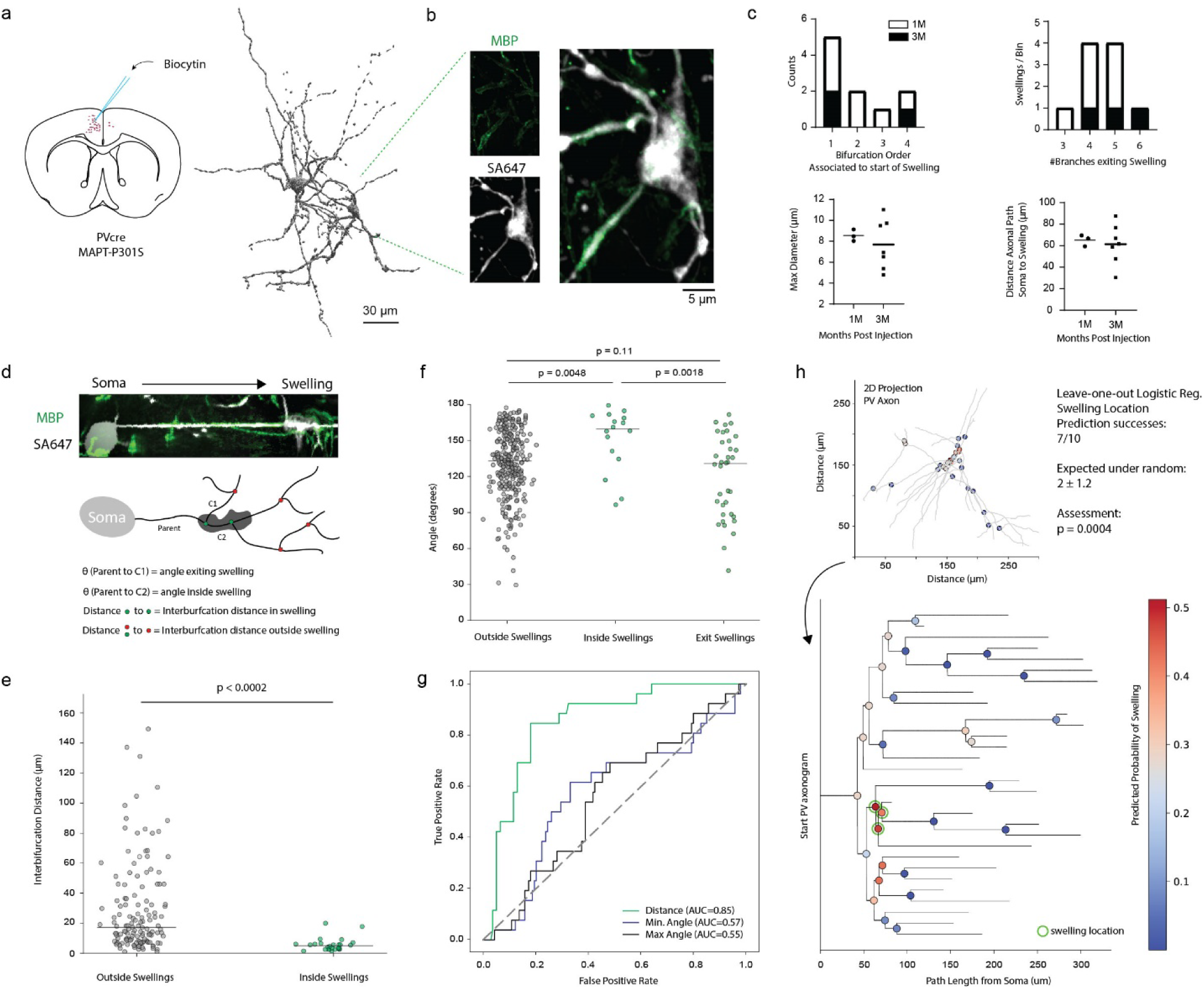
Dystrophy of axonal bifurcations can occur at different branch order points along the axonal arbor and their location can be predicted via local axon arbor geometry. **(a)** PV neurons expressing MAPT-P301S for 1 or 3 months were filled with biocytin using glass capillary and whole cell access prior to fixation and staining with Streptavidin Alexa647. Right showing a filled PV neuron with soma and swelling visible **(b)** Axonal dystrophy found in the neuron shown in (a), note MBP positivity of 2 emanating branches. **(c)** Descriptive parameters of the identified swellings, including bifurcation order presenting with swelling, number of branches exiting swelling, maximum swelling width, and length along the axonal path at which swelling started (n = 10 axonograms, N=5 mice). **(d)** (upper) Example of a biotin filled and streptavidin-647 stained PV neuron, that is digitally straightened along its axon from the soma to the bifurcation associated swelling, and (lower) schematic explaining the measurement of parent-child angles and inter-bifurcation distances. **(e)** Measured inter-bifurcation distances outside of swellings and inside of swellings, within-neuron permutation test, p < 0.0002, black line showing median. **(f)** Parent-child angles for branch-pairs outside of the swellings, inside swellings, and those exiting swelling. Within-neuron permutation tests, Holm-corrected, inside vs exiting, p = 0.0018; inside vs outside, p = 0.0048; exiting vs outside, p = 0.11, black line showing median. **(g)** ROC curve for logistic regression model to assess predictive power of inter-bifurcation distance (green), minimum bifurcation angle (blue), and maximal bifurcation branch pair bifurcation angle (black). Leave-one-neuron-out AUC: distance 0.852 (average precision 0.44, base rate 0.158), minimum angle 0.574, maximum angle 0.555; distance exceeded chance under within-neuron label permutation (null mean 0.41; p < 0.0002, 5,000 permutations). **(h)** 2D projection of a reconstructed Parvalbumin axon (upper) schematized as an ‘axonogram’ below, with color scale from blue to red matching the predicted probability of a swelling occurring as assessed by a leave-one-out based logistic regression model based on inter-bifurcation distance. Actual swelling location indicated by green circles. Top-1 p = 0.0004, top-3 p = 0.0297 (exact Poisson-binomial, one-sided).

Across the reconstructed axons, the swellings occurred at a range of axonal branch orders, from the first bifurcation to higher-order bifurcations, indicating that vulnerability is not confined to a specific depth within the arbor (Fig.4c). All reconstructed dystrophies included bifurcations, as denoted by ≥ 3 axonal branches emanating from the swelling (Fig.4c). To assess for potential geometric risk factors underlying the focal swellings in PV⁺ axons, we developed an axon-structure analysis pipeline. For each reconstructed axonal tree we extracted the angles formed between parent and child branches and the path-length distances separating consecutive bifurcations (Fig.4d). This approach allowed direct comparison between “swelling” and “non-swelling” bifurcations within and across the reconstructed arbors, as well as the construction and testing of a simple predictive model of local vulnerability. Geometric context revealed a clear signature of swelling localization. Bifurcations falling within swellings were separated by markedly shorter inter-bifurcation distances than non-swelling bifurcations (Fig. 4e). Thus, swellings tend to form in densely branching regions, where consecutive bifurcations occur within short distances. Parent-child angles within the swelling were straighter than at non-swelling bifurcations or at branches exiting the swelling, whereas exiting branches did not themselves differ from non-swelling bifurcations (Fig. 4f).

To evaluate whether the geometric features can predict swelling sites, we tested logistic regression models with a leave-one-out (LOO) cross-validation scheme. Here, one neuron is held out for testing while models are trained on the remaining neurons, and the process is repeated across all neurons. We trained logistic models on minimum and maximum parent-child angle and inter-bifurcation distance (Fig. 4g). The inter-bifurcation distance model performed strongly (LOO AUC 0.852) and exceeded chance under a permutation test respecting the clustering of bifurcations within neurons. Performance was unchanged when whole animals rather than individual neurons were held out in training sets (Leave-one-animal-out AUC for distance, 0.852). Minimum and maximum parent-child angles showed only weak predictive performance (LOO AUC 0.574 and 0.555, respectively), and neither improved performance when combined with distance (ΔAUC = - 0.0086 & −0.0003, Holm-adjusted permutation p = 0.780 & 0.508, for respective additive models).

We next asked whether the distance model could be used to predict locations of swellings in individual axonal arbors. To test this possibility, we generated bifurcation risk maps using the LOO inter-bifurcation distance model (Sup.Fig.6). With this approach, the top-ranked bifurcation matched a true swelling location in 7/10 neurons and appeared within the top three in 8/10, exceeding a Poisson-binomial null (top-1 p = 0.0004; top-3 p = 0.0297, Fig.4h). Performance was unchanged when the model was trained excluding all neurons from the same animal. Together, these findings reveal a distinct geometric signature associated with neurodegenerative risk in PV⁺ axons: short inter-bifurcation distances constitute the primary geometric correlate of focal dystrophy, with swellings preferentially located along relatively straight parent segments embedded within densely branched regions.

### Swellings of axonal bifurcations are structurally dynamic sites of axon severing and formation of myelin dystrophy

Because bifurcation-associated swellings emerged as focal points of pathology, we next investigated whether these structures mark sites of active axonal severing rather than stable morphological abnormalities. To bring into view the temporal relationship between formation of axonal swellings and eventual axon loss, we employed chronic *in vivo* two-photon imaging of P301S PV neurons, repeatedly imaging the same swellings at one-week intervals (Fig.5a). A total of 48 swellings were tracked longitudinally. To visualize potential evolution of oligodendrocytic dystrophy associated to the dystrophies, we co-injected a virus carrying constructs for membrane targeted-eGFP under MBP promoter. This effectively labeled the adjoining processes and core of a subset of the imaged swellings, where the PV neuron and oligodendrocyte responsible for its myelination were both successfully transduced.

**Figure 5.**
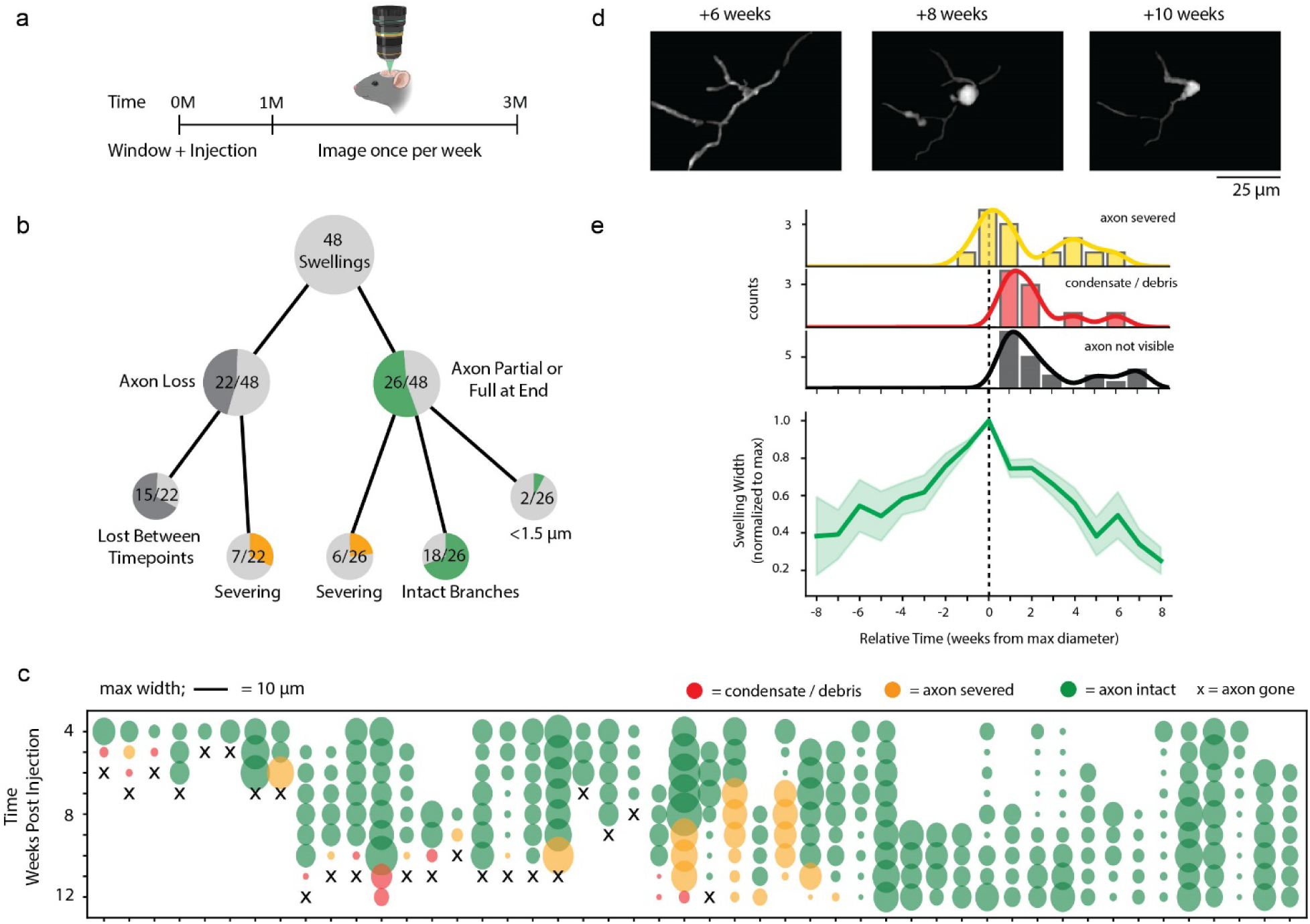
Dystrophies of axonal bifurcations are structurally dynamic and sites of axon severing. **(a)** Mice expressing the Tauopathy construct and eGFP under MBP promoter were longitudinally imaged at 1-week intervals to track individual swellings (N=3 mice). **(b)** Fate tree of the 48 imaged swellings. **(c)** Maximum swelling width across timepoints, further color coded with yellow indicating axon severed (partial loss of axon arbor), red to indicate axon lost but remnant debris/condensate visible, and x denoting complete loss of imaged axon. **(d)** 2D projection of a segmented imaging volume showing a PV axon with a bifurcation associated swelling that increases in size, before losing an axonal segment. **(e)** Swelling diameter and annotated events aligned to timepoint with maximum swelling cross-section.

The dataset revealed that bifurcation-associated swellings are structurally dynamic (Sup.Fig.7, example time series). Tracking the maximal width of each dystrophy showed trajectories that either progressively increased in diameter or fluctuated between growth and partial regression across weeks (Fig.5c). We annotated three outcomes: axon severed (‘yellow’, loss of one or more branches emerging from the swelling), condensate formation (‘red’, only a small fluorescent spot remaining without visible branches), and complete axon loss (‘x’, disappearance of the imaged axon). Across all 48 swellings, 30 (62.5%) acquired at least one of these fates within the imaging window. In 5/18 of the unresolved cases, the swelling reached its maximal diameter only at the final imaging session. Overall, 22/48 swellings (45.8%) culminated in complete axon loss during the observation period (Fig.5b,c). We frequently observed a residual condensate, a small fluorescent remnant localized to the former swelling site prior to complete loss. A reduction in the number of axonal branches emanating from the dystrophic region, consistent with local axon severing, was observed in 13/48 swellings (27.1%), and was present in each imaged subject (Fig. 5d).

To better understand the temporal relationship between swelling dynamics and axon fate, we aligned imaging sessions to the time point of maximal swelling diameter (Fig. 5e). By definition, condensate-only observations and complete axon disappearance occurred after the maximal measurable diameter had been reached. However, the time interval between peak swelling and subsequent axon loss varied across lesions, indicating heterogeneous progression after maximal expansion. First observations of severed, incomplete axons were distributed around the peak, with 1/13 detected before, 4/13 at, and 8/13 after maximal swelling diameter. Thus, bifurcation-associated swellings are dynamic sites of local branch severing, with progression to complete axon disappearance occurring at variable rates across imaged swellings.

Given our observation that swellings frequently arose within myelinated axon segments (Fig.3d), we next examined oligodendrocytes in the same fields of view. As with the neural component, oligodendrocyte processes adjoining the axonal unit showed highly dynamic responses to Tau-induced axon swellings. Paralleling the data presented above via IHF, oligodendrocyte processes enveloping the axon swelled together with the axon (Sup.Fig.8). In the majority of cases, the oligodendrocyte morphology corresponded closely to the dystrophic axon diameter. However, we also observed several instances of transient oligodendrocyte reactivity that did not match the physical dimensions of the axon dystrophy. Instead, at single timepoints, the oligodendrocyte membrane presented with an expanded diameter, which returned to baseline by the next imaging timepoint (Sup.Fig.9). This suggests that while oligodendrocyte membranes initially stretch to match the increasing dimensions of the underlying axonal swelling, the glial unit may experience a downstream set of dystrophic signaling and structural responses during the progression of the axonal dystrophy.

### Dystrophies of PV axonal bifurcations are present in cortex of human tauopathy patients

To check for the preservation of the observed phenotype across species, we returned to human CBD cases to assess for the presence of the bifurcation associated dystrophies in PV axons. To this end, we employed IHF in 100 µm thick free floating tissue sections. We found multiple examples of PV bifurcation-associated dystrophies in cortex of CBD subjects (Fig.6). Twenty-two such dystrophies were imaged using Airyscan super-resolution microscopy across four CBD donors. In 19/22 dystrophies, AT8 positivity was identified in emanating branches. At the swelling core, ten swellings had abundant Tau accumulation, nine showed discernible levels, whereas no AT8 was found in the core of three of the swellings. Around 40% (9/22) presented with MBP dystrophy (Fig.6a, Sup.Fig.10), whereas 13 did not. The presence of bifurcation-associated swellings with and without overt MBP dystrophy is consistent with the view that axonal geometry contributes to lesion localization, while overt myelin abnormalities may reflect an accompanying or secondary feature in a subset of lesions. As in the mouse model, the dystrophies had varying amounts of emanating branches, ranging from three to six (Fig.6b). Their maximum widths ranged from 2.8 up to 13.3 microns in length. The bifurcations within the swellings presented with short inter-bifurcation distances, corroborating the geometric vulnerability in the axonal arbor (Fig.6b-d). Thus, human CBD cortex recapitulates the bifurcation-centered, geometry-linked PV axonal dystrophies observed in our mouse model, consistent with a shared structural vulnerability across species.

**Figure 6.**
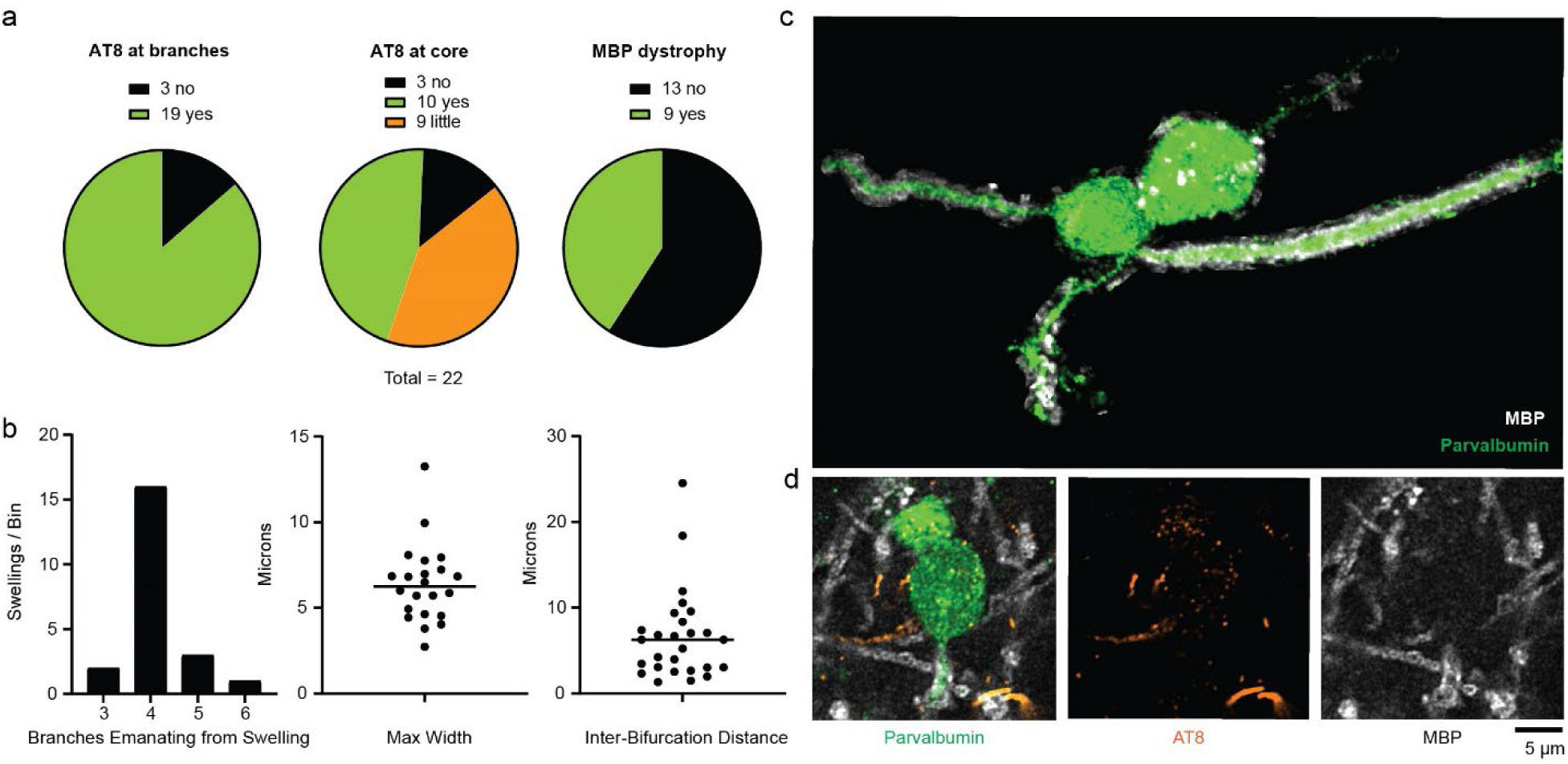
Dystrophies of PV axonal bifurcations are present in cortex of CBD patients. **(a)** Fraction of bifurcation associated swellings with AT8 neuropil thread at emanating branches, with AT8 at core and with MBP dystrophy. **(b)** Histogram of the number of branches emanating swellings, (center) observed maximum width of the axonal bifurcation associated dystrophies, and (right) measurements of the inter-bifurcation distances of the imaged swellings. **(c)** Example swelling from CBD cortex with AT8 positivity at core and no overt myelin dystrophy. Image shows the 3D segmented and masked axon dystrophy with 4 exiting branches. **(d)** A cropped single plane image showing the branch exiting the dystrophy to the left in panel C. Note AT8 positivity at the swelling core. Parvalbumin protein in green, AT8 in orange, and MBP in white.

## Discussion

Our results show that in both human primary tauopathy and a cell type restricted mouse model, parvalbumin neuron axons develop focal dystrophic swellings that arise at axonal bifurcations. These swellings accumulate organelles, precede local axon severing, and are associated with subsequent neuronal loss. Quantitative modeling demonstrates their enrichment at branch points and identifies local axonal geometry, especially short inter-bifurcation distances, as a key correlate of vulnerability. Together these findings indicate that vulnerability to Tau-induced degeneration within the PV axonal arbor is spatially patterned, and that the sites at risk can be identified from local geometry alone.

Our data support a model in which neuritic dystrophy is an early structural feature of Tau-induced PV neuron degeneration. *In vivo*, neuritic swellings were already present at the first imaging time point and increased in prevalence over the same period in which PV soma loss emerged. At the population level, swelling burden within imaged volumes coincided with subsequent soma loss most clearly during the later 2M-3M interval. This correlation should not be overinterpreted as a direct neuron-by-neuron fate relationship, as the imaging conditions and dense PV neuritic labeling in this longitudinal dataset did not permit reliable assignment of individual swellings to their originating somas. Nevertheless, the interpretation that axonal pathology precedes soma loss is supported by convergent observations: *ex vivo* IHF as well as biocytin-filled patch-clamp reconstructions identified focal axonal swellings in neurons with morphologically intact somas, and longitudinal tracking of individual bifurcation-associated swellings showed progression to local branch severing and axon disappearance. Together, these findings suggest a dying-back-like process in which the PV axonal arbor, and particularly its branch points, represent early sites of structural failure during Tau-induced degeneration.

Little is known about the biology of axon bifurcations in the adult brain, but developmental and ultrastructural work indicate that they are specialized microdomains (*17*). Axon collateral branches often emerge from discrete actin “patches” along the shaft that depend on Arp2/3-mediated actin nucleation, and disruption of this machinery reduces branch formation (*18–21*). Microtubule-severing proteins such as spastin and katanin generate short microtubule fragments that invade nascent branches, highlighting localized cytoskeletal remodeling at branching sites (*22*, *23*).

Ultrastructural and organelle mapping studies further show that mitochondria, ribosomes, endoplasmic reticulum and presynaptic vesicles are enriched at or near branch points, where they are thought to support local energy demands, calcium handling and protein synthesis (*17*, *23–25*). At the same time, reviews highlight that the long-term maintenance, trafficking properties and vulnerability of established branches in the adult brain, particularly in vivo, remain poorly understood (*19*, *20*, *23*). Our data extend this picture in two ways. First, they provide *in vivo* evidence that mature axon branch points carry a distinct structural vulnerability under pathological conditions. Second, and more specifically, they identify short inter-bifurcation distance, a property of local branching topology rather than of any individual bifurcation, as the primary geometric correlate of that vulnerability. This suggests that it is not simply the presence of a branch point, but the density of branching within a local axonal domain, that constitutes the strongest risk factor for tau-induced branch-point dystrophy.

The available evidence suggests that axonal bifurcations act as microdomains with complex trafficking logistics. Under pathological Tau, which can perturb microtubule organization and stability, hinder kinesin-1 interaction with microtubules and impair mitochondrial energetics (*9–11*), these junctions may become local bottlenecks for axoplasmic flow. Consistent with this idea, transport models in branched neurons show that local microtubule disruption can produce focal cargo accumulation and uneven delivery to downstream branches (*26*), paralleling the organelle and motor-protein accumulation observed here. A downstream consequence could be functional isolation of distal segments that now receive inadequate anterograde supply and retrograde clearance, predisposing them to degeneration. This sequence is consistent with occasional observations in our longitudinal imaging of asymmetric branch loss at swelling sites, and parallels the one-sided neurite failure and marked structural plasticity documented around amyloid plaques (*27*).

Neuropil threads are a defining histopathological feature of tauopathies, yet their cellular origin is often difficult to resolve because affected neurites are embedded in dense neuropil and disconnected from identifiable somas. By combining PV labeling with AT8 and MBP in thick human tissue, our data attribute tau-positive neuritic pathology, including myelinated axonal segments, to PV interneurons. This provides a framework for dissecting neuropil-thread burden by neuronal subtype in tauopathies. Our current study focused on PV neurites. It is, however, abundantly evident that a majority of neuropil Tau load observed in the analyzed CBD subjects lies in other cell types (Sup. Fig. 1). This raises two immediate questions: which additional neuronal subtypes contribute to neuropil load in CBD, and do these other unidentified neuronal subtypes present with similar sensitivity of bifurcations? While a subset of neurons can be identified based on characteristic expression profiles of subtype-defining proteins, many are defined by molecular compositions present primarily at their soma. We foresee that, in order to unambiguously elucidate the full range of afflicted neurons contributing to neuropil load, large-scale high-resolution imaging in multiplexed IHF experiments or spatial transcriptomics preparations will be required, enabling tracing of neurites back to their somatic origin. The workflow established here, and further improvements thereof, could equally be deployed in additional tauopathies to delineate neuropil load contribution by cell type.

Several limitations of the current study merit discussion. The AAV-mediated P301S expression system, while enabling cell-type restricted interrogation of tau-induced pathology, represents an accelerated and cell-autonomous model that likely exceeds the Tau burden and tempo of endogenous tauopathy. The degree of soma loss observed here should therefore be interpreted accordingly, rather than as a direct prediction of neuronal loss rates in human disease. Additionally, while the geometric analysis identifies short inter-bifurcation distance as the primary structural correlate of swelling localization, the molecular mechanisms that render densely branched axonal regions specifically vulnerable, whether related to local cytoskeletal organization, mitochondrial distribution, trafficking logistics, or further sources, remain to be defined. Future studies combining ultrastructural analysis and live transport imaging at branch points will be necessary to resolve these questions.

In sum, the convergence of findings across *in vivo* imaging, quantitative modeling, and human postmortem tissue identifies bifurcation-associated dystrophies as a recurrent locus of PV axonal pathology in tauopathy. Beyond the specific context of PV interneurons, these findings raise the broader possibility that axonal branching topology corresponds with degenerative vulnerability across neuronal subtypes, and that geometry-linked compartmental failure may represent a general principle of tauopathy progression. Conversely, absence of such a phenotype in other neuronal subtypes would indicate a locus of selective vulnerability within a molecularly defined population.

## Materials and Methods

### Ethics

All animal experiments were conducted in accordance with animal welfare regulations of the Government of Upper Bavaria (approval no. Vet_02-20-207). Human brain tissue was provided by the Neurobiobank Munich under a biobank protocol approved by the Ethics Committee of LMU Munich (approval no. 345-13). Donors provided written informed consent during life. Use tissue for the present study was approved by the Ethics Committee of LMU Munich (approval no. 22-0729) and complied with the Declaration of Helsinki and its later amendments.

### Virus generation

The complete plasmid sequence of AAV-EF1A-DIO-MAPT(P301S)-P2A-mKate2-WPRE and AAV-EF1A-DIO-mKate2-WPRE utilized for AAV5 production is available in supplementary methods. AAV5 production was carried out by VectorBuilder using the triple transfection-based method in HEK293-AAV followed by purification using cesium chloride (CsCl) gradient ultracentrifugation (VectorBuilder). The concentration of genome-containing AAV particles was determined by qPCR. rAAV2.1-MBP-EGFP particles were produced according to Challis et al., (2019) (*28*). Briefly, HEK293T cells (ATCC crl-3216) were transfected using polyethylenimine (PEI) with a 1:4:2 molar ratio of helper (pAdDeltaF6; Addgene # 112867), capsid (pAAV2/1; Addgene # 112862) and the plasmids of interest. Supernatant was harvested at 72h and 120h after transfection and kept at 4°C and cells were collected at 120h. The supernatant, combined with PEG solution overnight, was centrifuged at the speed of 4000 g for 30 minutes at 4 °C and the pellet was mixed with the cell lysate and loaded onto an iodoxanol gradient. Centrifugation was performed at 350,000 g for 2 hours and 25 minutes at 18 °C with slow acceleration, before virus was loaded on Amicon filter and washed several times. Titers were determined using qPCR. Viral aliquots were stored at −80°C until use.

### Animals

Mice, both male and female were used and held on a 12-h light/dark cycle with food and water ad libitum. Two strains used were the PV-Cre (B6;129P2-Pvalbtm1(cre)Arbr/J) and crossing of MitoTag mice (B6N.Cg-Gt(ROSA)26Sortm1(CAG-EGFP*)Thm/J) with PV-Cre mice to visualize mitochondria within PV neurites. The MitoTag mouse line was obtained from the group of Prof. Thomas Misgeld. Mice were genotyped via ear punches with PCR. Primers used for Cre line were Cre-F AGC CTG TTT TGC ACG TTC ACC with Cre-R GGT TTC CCG CAG AAC CTG AA. Primers used for Rosa26-MitoTag-OMM were Mito-1 GCA CTT GTC CTC CCA AAG TG with Mito-2 CAT AGT CTA ACT CGC GAC ACT G and Mito-3 CAA GAT CCG CCA CAA CAT CG with Mito-4 TAT CTC ACG AAG GCC CAA AC.

### Virus Injection and Chronic cortical cranial window implantation surgeries

PV-Cre were injected with either a vector inducing tauopathy (AAV-EF1A-DIO-MAPT(P301S)-P2A-mKate2-WPRE) or a control construct (AAV-EF1A-DIO-mKate2-WPRE), or a 1 to 1 combination of tauopathy vector with the Oligodendrocytic eGFP vector (AAV concentrations, 10^13^ vg/ml all preps). Cortical cranial windows were fitted in the same surgery. Mice were anesthetized with a mixture of Medetomidin, Midazolam and Fentanyl at 0.5, 5 and 0.05 mg/kg bodyweight respectively, and received 10 mg/kg Dexamethason to reduce inflammatory responses prior to placement on stereotactic frame. Following skin excision and periosteum removal, the exposed skull was lightly scored with a scalpel and an initial mildly corrosive adhesive layer was applied and activated (IBond Self Etch, Kulzer 66046243). A 4mm craniotomy centered on the secondary motor cortex (relative bregma A.P.+1.0 mm M.L.+1.2mm) was drilled using a Neurostar automated drilling system until the skull disk could be manually removed. The dura mater was removed manually. Viral injection followed the craniotomy (∼A.P.+1.0 mm M.L.+1.2mm, depth 300µm from pia). For all conditions the injected volume was 600nl at an injection rate of 60nL/min. After the virus injection the craniotomy area was cleaned with PBS and a 4mm circular glass cover slip was fitted and fixed in place with tissue adhesive glue (Surgibond tissue adhesive, Praxisdienst, 190740). The exposed skull was subsequently covered with dental cement (Gradia Direct Flo BW, Spree Dental, 2485494), a headbar with a central hole was placed over the window, and the adhesive was cured with UV light. After surgery mice received 5 mg/kg Enrofloxacin as an antibiotic, 25 mg/kg Carprofen to reduce inflammation and 0.1 mg/kg Buprenorphin as an analgesic. A mixture of Atipamezol and Flumazenil (2.5 and 0.5 mg/kg) was used to antagonize the anaesthesia. For experiments in which only a virus injection was employed, the same medications and anaesthesia were administered, and injections were carried out via a small bur hole located at relative to bregma at A.P.+1.0 mm M.L.+1.2mm and at an approximate depth of 300µm from pia.

### Immunostaining

Anesthetized mice were transcardially perfused with phosphate-buffered saline (PBS), followed by 4% paraformaldehyde (PFA). After a 16h postfixation period at 4°C in 4% PFA brains were stored in PBS with 0.05% sodium azide (NaN_3_). Coronal brain slices of 50 µm thickness were obtained by cutting with a vibratome (VT1200S, Leica Biosystems). Long term formalin fixed human tissues were transferred to PBS with 0.05% sodium azide, stored at 4° C and sliced at 100um thickness. Human sections underwent antigen retrieval in a pH 6 sodium citrate based solution for 1hr at 100 degrees Celsius, and autofluorescence was quenched and bleached using broad spectrum white lights with slices immersed in 1× PBS with 4.5% H₂O₂ and 20mM NaOH for 1h. All stains were performed on free-floating sections. Slices were first permeabilized with 2% Triton for 2 hours and subsequently blocked with blocking solution (10 % normal goat serum and/or 10 % normal donkey serum in 0.3 %Triton and PBS) for 2 hours at RT. Primary antibodies were incubated over-night at 4°C, followed by washing and secondary antibody incubation for 1 hour at room temperature, protected against light. Slices were mounted with mounting medium, containing DAPI (RotiMount FluorCare, Roth). Primary antibodies used were: rabbit anti-PV (1:1000, Abcam, AB11427), mouse-AT8 (1:500, ThermoFisher, MN1020), rat anti-MBP (1:500, Abcam, AB7349), guinea-pig anti-tagRFP/mKate2 (1:1000, Cancer Tools, 155267), guinea-pig anti-PV (1:1000, Sysy, 195 004), rabbit-T22 (1:1000, Merck, ABN454), rabbit-pS396 (1:1000, Invitrogen, 44-752G), rabbit pT231 (1:1000, Abcam, AB151559), rabbit-pS404 (1:1000, Abcam, AB92676), rabbit anti-LAMP1 (1:500, Abcam, AB24170), chicken anti-GFP (1:10.000, Abcam, AB13970), rabbit anti-Kif5a (1:1000, Proteintech, 84013-5-RR). Secondary antibodies used were: goat-anti-gp-555 (Invitrogen, A21435), goat-anti-gp-488 (Invitrogen, A11073) goat-anti-ch-488 (Invitrogen A11039), goat-anti-rb-647 (Invitrogen A21245), goat-anti-rat-647 (Invitrogen A21247), goat-anti-ms-647 (Invitrogen A31571), donkey-anti-ms-555 (Invitrogen A32773), goat-anti-rat 488 (Thermo, A-11006). Images were acquired with a Zeiss LSM900 airyscan microscope (Carl Zeiss).

### Quantification AT8 and MBP in human PV neurites

Images were acquired on a Zeiss LSM900 airyscan microscope, with voxel sampling size of 43×43×150 nm (xyz). SWC traces along PV neurites were generated with the Simple Neurite Trace (SNT) ImageJ-plugin. The paths were smoothed, and straightened to sample cross-sections orthogonal to each neurite’s path. A mean projection across 3 z-slice steps centered on the traced path position (0.15um step size, 0.45um projection) was generated at each path position. At each cross-section along a neurite, AT8 or MBP positivity was manually annotated. Total neurite length and the lengths of AT8+, MBP+, and double-positive neurites were subsequently aggregated at the subject levels.

### Probability Swellings Occur at Branch Points

To assess whether the association of neuritic swellings with axonal bifurcations exceeded chance levels, publicly available PV axon reconstructions (*16*) were linearized at 0.1 µm resolution, with branch points annotated. PV bifurcation associated Swelling lengths along the axon were determined empirically in 3- and 6-month post-injection mice. Swelling length was defined as the longest path through a given swelling, with the swelling being the portion of the axon in which diameter exceeded 3x FWHM of mKate2 signal along the neurite path along a section of the neurite far removed from the swelling. A sliding-window analysis was implemented in which virtual windows, matched to empirically measured swelling lengths, were moved along each reconstructed axon in non-overlapping steps to estimate the probability that a randomly positioned axonal segment would include a branch point. All trajectories from the soma to terminal axonal points were included per neuron. Because each soma-to-terminal trajectory was traced independently, proximal segments were represented multiple times across paths. This approach preserves the arbor topology and slightly overweighs regions near bifurcations, rendering the baseline probability estimate conservative. The estimated baseline probability was then compared to the observed frequency of bifurcation-associated swellings ex vivo using a one-sided binomial test. Confidence intervals for observed proportions were calculated using the Wilson method with a two-sided 95% confidence level, and robustness of the enrichment was evaluated through bootstrapping with replacement across neurons available in the public dataset. The analysis was further repeated across a range of window lengths to examine sensitivity to swelling size, encompassing and extending beyond the empirically determined swelling length distribution.

### Quantification accumulations at dystrophies

Images were acquired on a Zeiss LSM900 airyscan microscope, with voxel size of either 43×43×170 nm or 50×50×190 (xyz). Neurite centerlines through swellings were manually traced via SNT plugin image-j plugin. Image stacks were straightened along the traced path (single z-slice) and interpolated at 1-pixel intervals. Fluorescence intensity profiles were extracted along the traced neurite axis by averaging signal across a central orthogonal bin (0.25 µm). Profiles were aligned to the position of maximal swelling diameter, and binned into 100 bins across the common overlapping distance range and plotted as mean ± SEM. ΔF/F was calculated per trace using the mean fluorescence between −25 and −15 µm relative to the swelling center as baseline. Mean fluorescence values in baseline (−25 to −15 µm) and peak (−5 to +5 µm) regions were compared per swelling using paired two-tailed t-tests or Wilcoxon signed-rank tests, following assessment of normality.

### Longitudinal in vivo 2P Microscopy

The two-photon microscope system was the Femtonics ATLAS3D fitted with a Coherent Fidelity (1040nm), and a Coherent Chameleon Ultra-II tunable laser set at 920nm, to excite mKate2 and eGFP respectively. Laser output was attenuated to <50mW average power at the focal plane. 3 weeks after surgery mice were gradually accustomed to head-restraint and placement on the running wheel used for in vivo imaging through multiple training sessions. In vivo imaging experiments began 1 month after cranial window implantation. *Experiment 1:* For experiments tracking somatic fate, mice were imaged at one-month intervals. Somas identified in the first imaging session were manually tracked along imaging sessions. The experiment was conducted in 2 cohorts (N=3 per group for each cohort). Imaging volumes were aligned to pia. For cohort 1, 1 region per mouse was imaged (274×274×300 µm, 0.39 µm/pixel, 0.6 µm/z-step), For cohort 2, 3 regions per mouse were imaged (300×300×300 µm, 0.37 µm/pixel, 1 µm/z-step); volumes lost due to decreased cranial window clarity between timepoints (1 Volume, P301S group), and volumes with percent soma loss exceeding group-wise third quartile by more than 1.5 × the P301S interquartile range (Tukey’s rule; 2 volumes, P301S group) were excluded from further analysis, resulting in final N_p301s_=6, volumes = 9; N_control_ = 6, volumes=12. Where multiple imaging volumes were acquired from the same mouse, measurements were averaged across volumes at each timepoint or longitudinal interval prior to statistical analysis. For identification of swellings in corresponding volumes, swellings were defined as segments of neurites in which the diameter was >3x mean of healthy neurites. For longitudinal analyses of PV soma survival and neuritic swelling burden, each mouse was summarized by the mean of its measurements at months 2 and 3 minus its own month 1 baseline. For survival, month 1 was fixed at 100% by definition. Per-mouse changes from baseline were compared between control and P301S groups using two-sided exact permutation Mann-Whitney tests. The baseline diameter of healthy neurites was determined by measuring diameters of 50 healthy neurites from the controls group (3 volumes from 3 separate subjects, 15-20 neurites per volume). The relationship between swelling burden at timepoint N and PV soma loss between timepoints N and N+1 was assessed separately for each longitudinal interval using one-sided Spearman rank correlations, based on the directional hypothesis that greater swelling burden would be associated with greater subsequent soma loss. *Experiment 2:* For experiments tracking swelling longitudinally, mice were imaged at one-week intervals. Images centered on a bifurcation were 75 by 75 by 40 microns, pixel width/height 0.094 microns, 0.3micron step size Z. A subset of dystrophies included in the analysis appeared in the imaged volumes subsequent to start of imaging.

### Identification and measurement of human PV axonal dystrophies

IHF triple stained sections of CBD cortical tissue (AT8, PV, MBP) were screened and bifurcation associated dystrophies were identified qualitatively as overt lesions showing a clear, abrupt enlargement of neurite diameter. Maximum widths and inter-bifurcation distances were subsequently measured on the identified dystrophies.

### Patch-clamp for Axonal Reconstructions and Electrophysiology

Slice preparation; 1 and 3 months after viral injection, male and female mice were anesthetized with isoflurane, transcardially perfused, decapitated, and brains were extracted. Coronal sections containing the secondary motor cortex (250μm) were prepared on a vibratome (Campden 7000smz2) in Glycerol slicing solution containing (in mM) 250 Glycerol, 2.5 KCl, 1.2 NaH_2_PO_4_, 26 NaHCO_3_, 10 HEPES, 5 MgCl_2_, 1 CaCl_2_, 11 Glucose carbogenated with 5% CO_2_/95% O_2_. Slices were incubated in ACSF containing (in mM) 125 NaCl, 2.5 KCl, 1.2 NaH_2_PO_4_, 26 NaHCO_3_, 10 HEPES, 2 MgCl_2_, 2 CaCl_2_, 11 Glucose carbogenated with 5% CO_2_/95% O_2_ at 35C for 30min and then allowed to recover at room temperature for 30min. After recovery, slices were placed in the recording chamber and perfused with ACSF at a rate of 1.5-2ml/min at 30-32C. All recordings were performed within 6h of slice recovery. Whole-cell patch clamp; PV neurons in layer II and III were visualized using infrared differential interference contrast video microscopy and expression of mKate2 via fluorescence colocalization (Olympus BX50WI; QImaging Retiga ELECTRO). Whole-cell current-clamp was performed using HEKA EPC 10 USB. Borosilicate glass pipette (Sutter Instrument) electrodes (4-6 MΩ) contained (in mM) 140 potassium gluconate, 5 KCl, 10 HEPES, 0.2 EGTA, 2 NaCl, 4 ATP, 0.5 GTP, 0.2% biocytin (pH 7.2-7.3, 280-290 mOsm). Resting membrane potential was recorded and averaged across 3 min. To investigate neuronal excitability, neurons were held at −70mV and depolarizing current steps (2s, 0-400pA in 50pA step). Cells had serial resistance < 25 MΩ, and input resistance > 100 MΩ. Data were digitized at 20k Hz. The calculated liquid junction potential of 14.6mV was compensated. Recorded slices were immersed overnight in 4% PFA for 3D morphological reconstruction of biocytin filled neurons. For overall input-output responses, four-parameter logistic curves were fitted separately at 1 and 3 months; group differences were assessed by neuron-level permutation of treatment labels (20,000 permutations), using the extra-sum-of-squares F statistic comparing a common curve with a model allowing group-specific half-maximal current and curve steepness, while sharing the lower and upper plateaus. Slices containing filled neurons were labeled overnight with MBP antibody (see below), before being stained with anti-rat-alexa488 antibody and streptavidin-alexa647. Axonograms were generated using Simple Neurite Trace (SNT) plugin for ImageJ (Fiji). Generated axonograms were exported as .swc for further analysis (see below).

### Axonogram Analysis and Bifurcation Risk Modeling

To assess whether local axonal geometry predicts the occurrence of focal swellings, we quantified bifurcation-level features from reconstructed PV neuron axonal arbors. For each bifurcation we extracted the inter-bifurcation distance, defined as the 3D path-length separating the bifurcation from its nearest upstream or downstream branch point, and the minimum and maximum parent-child angle, i.e. the smallest and largest angle between the parent segment and either child branch. Angles were measured between three points: the bifurcation itself, a point 10 µm retrograde along the parent, and a point 10 µm anterograde along the child (where the adjacent branch segment was shorter than 10 µm, its endpoint at the next branch point, terminal, or root was used), such that a straight continuation approaches 180°. Bifurcations for which any angle or the inter-bifurcation distance was undefined, for example where no further branch point occurred before the end of the reconstruction, were excluded, leaving 165 bifurcations. Each was labelled “swelling” or “non-swelling” according to the presence of a dystrophic enlargement; each neuron carried a single swelling, which could span several consecutive bifurcations. Binary logistic regression models were fit to predict swelling localization. Variants tested were an inter-bifurcation-distance-only, minimum-angle-only, maximum-angle-only, and additive minimum-angle or maximum angle plus-distance models. Performance was evaluated by leave-one-neuron-out cross-validation, in which each neuron was held out for testing while the model was trained on the remaining neurons, and summarized by ROC AUC and average precision. Average precision was interpreted relative to the proportion of bifurcations that were swelling associated, corresponding to the value expected from random sampling (0.158). The distance-only model was additionally evaluated using leave-one-animal-out cross-validation, in which all neurons from one animal were excluded from training and used as the test set for that animal. As each neuron carries one swelling which can span adjacent bifurcations, geometric features were compared by two-sided Mann-Whitney U statistics evaluated under permutation of swelling labels within each neuron, preserving the number of swelling-associated bifurcations per neuron. The same within-neuron permutation was used to test the cross-validated distance AUC against chance. To test whether either angle metric improved discrimination beyond distance, each angle variable was permuted while distance and swelling labels were held fixed; the two resulting ΔAUC tests were Holm-corrected. All permutation tests used 5,000 permutations with a fixed seed. Per-neuron risk maps were generated using the LOO distance-only model to assign swelling probabilities to all bifurcations. For each neuron, the top-1 and top-3 highest-probability bifurcations were tested against an exact Poisson binomial null derived from the number of bifurcations and swelling-associated bifurcations per neuron, giving one-sided probabilities for the observed tagging rates.

### Statistics

Statistical analyses were performed using Prism and Python. Detailed information on statistical analyses and experimental sample sizes is provided in the main text, figure legends, methods sections, and supplementary information. Unless otherwise stated, data are presented as mean ± SEM. Probability threshold for statistical significance was *p* < 0.05, and exact *p* values are reported throughout.

## Contributions

PF and JH conceptualized the study. PF performed experiments, analysis, and wrote the first draft of the manuscript. YZ performed patch clamp recordings and analysis. TN, NL, CV, LP contributed to experiments, data interpretation, project administration. LP designed neuron targeting constructs. FMB designed and prepared oligodendrocyte targeting constructs. All authors revised and approved the final manuscript.

## Funding

This project was funded by; the Munich Cluster of Systems Neurology (SyNergy; project ID EXC 2145 / ID 390857198), LMU Munich, Munich, Germany (to JH, FMB), German Center for Neurodegenerative Diseases (DZNE) Foundation, Innovative Minds Program (#01886) (to PF).

## Competing Interests

The authors declare no competing interests.

## Availability of data, code, and materials

All data supporting the findings of this study are available in the main text, supplementary information, and supplementary data file. Reconstruction files (.swc) and the analysis code for the bifurcation risk modelling are available at Zenodo (https://doi.org/10.5281/zenodo.22801634 & https://doi.org/10.5281/zenodo.22808315).

## Acknowledgments

The authors thank Dr. G. Mitteregger, Dr. M. Schneider, and the DZNE animal house team for excellent administrative support and animal care. We also thank Fang Zhang, and Dr. N. Buresch for dedicated technical support, and Dr. Sigrun Roeber for support from the NBM. We especially thank the donors and their families for making this research possible.

## Supplementary Materials

Supplementary Data File 1

Supplementary Figures 1 to 10

Supplementary Table 1

Supplementary Video 1 Vector Sequences

## Notes

### Competing Interest Statement

The authors have declared no competing interest.

