## Supplementary Material for "Axonal Branch Points of Parvalbumin Interneurons are Focal Sites of Tau-Induced Degeneration"

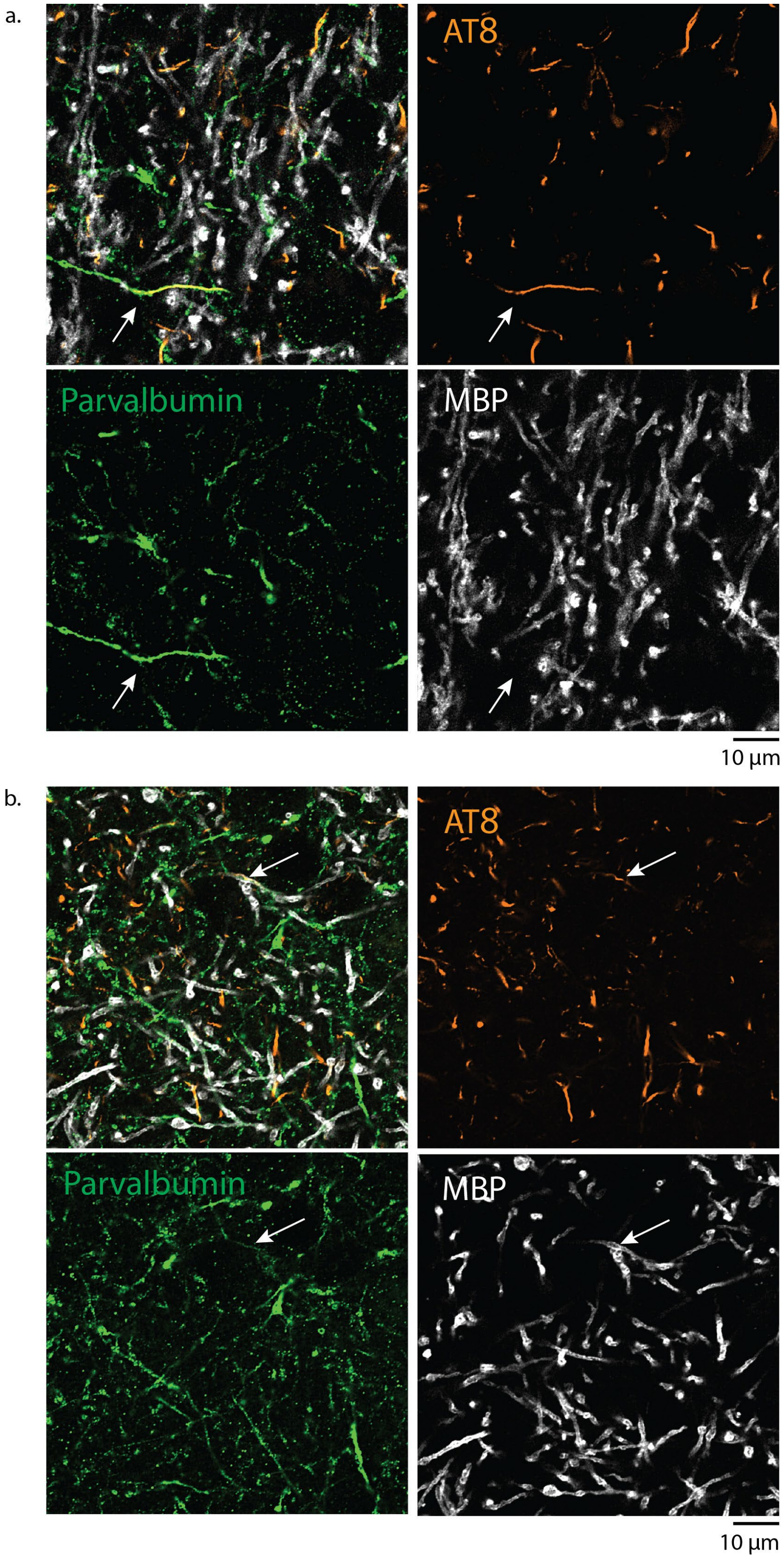


**Sup. Fig 1.** Single optical section of an Airyscan z-stack acquired from CBD donor tissue showing Parvalbumin (green), AT8 (orange), and MBP (white) IHF stain. **(a)** Neuropil thread in unmyelinated PV neurite, and **(b)** in a myelinated PV neurite, annotated by inset white arrow across color channels and merged image.

| Age | Diagnosis | Sex | Synuclein | Aß | TDP-43 | FUS | PMI (hrs) |
|---|---|---|---|---|---|---|---|
| 61 | CBD | Female | neg | neg | neg | neg | 28 |
| 52 | CBD | Female | neg | neg | neg | neg | 14 |
| 56 | CBD | Male | neg | neg | neg | neg | 44 |
| 56 | CBD | Female | neg | neg | neg | neg | 58 |

**Sup. Table 1.** Demographic and diagnostic information of CBD donors. All donors met neuropathological criteria for CBD and lacked significant Alzheimer's disease, synucleinopathy, or TDP-43/FUS pathology.

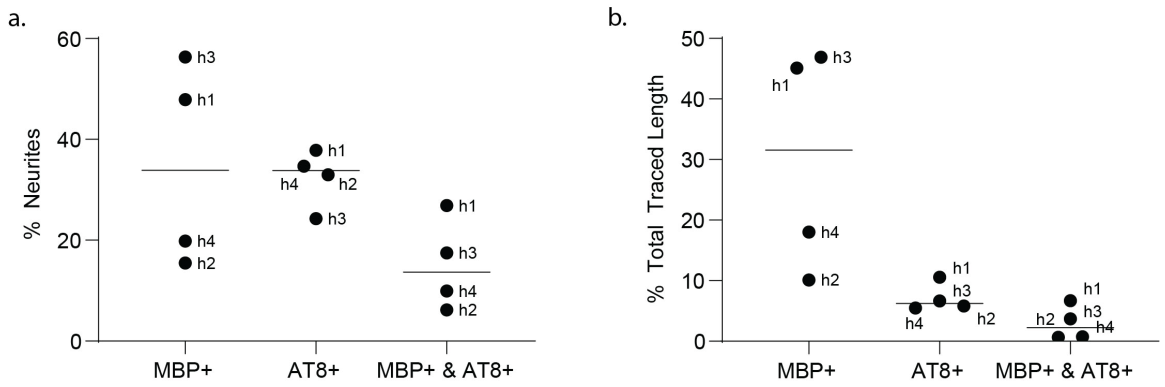


**Sup. Fig 2.** Quantification of MBP and AT8 overlap in traced PV neurites from CBD patient cortex. **(a)** Percentage of traced PV neurites classified as MBP-positive, AT8-positive, or double-positive in each case. **(b)** Percentage of total traced PV neurite length positive for MBP, AT8, or both markers in each case.

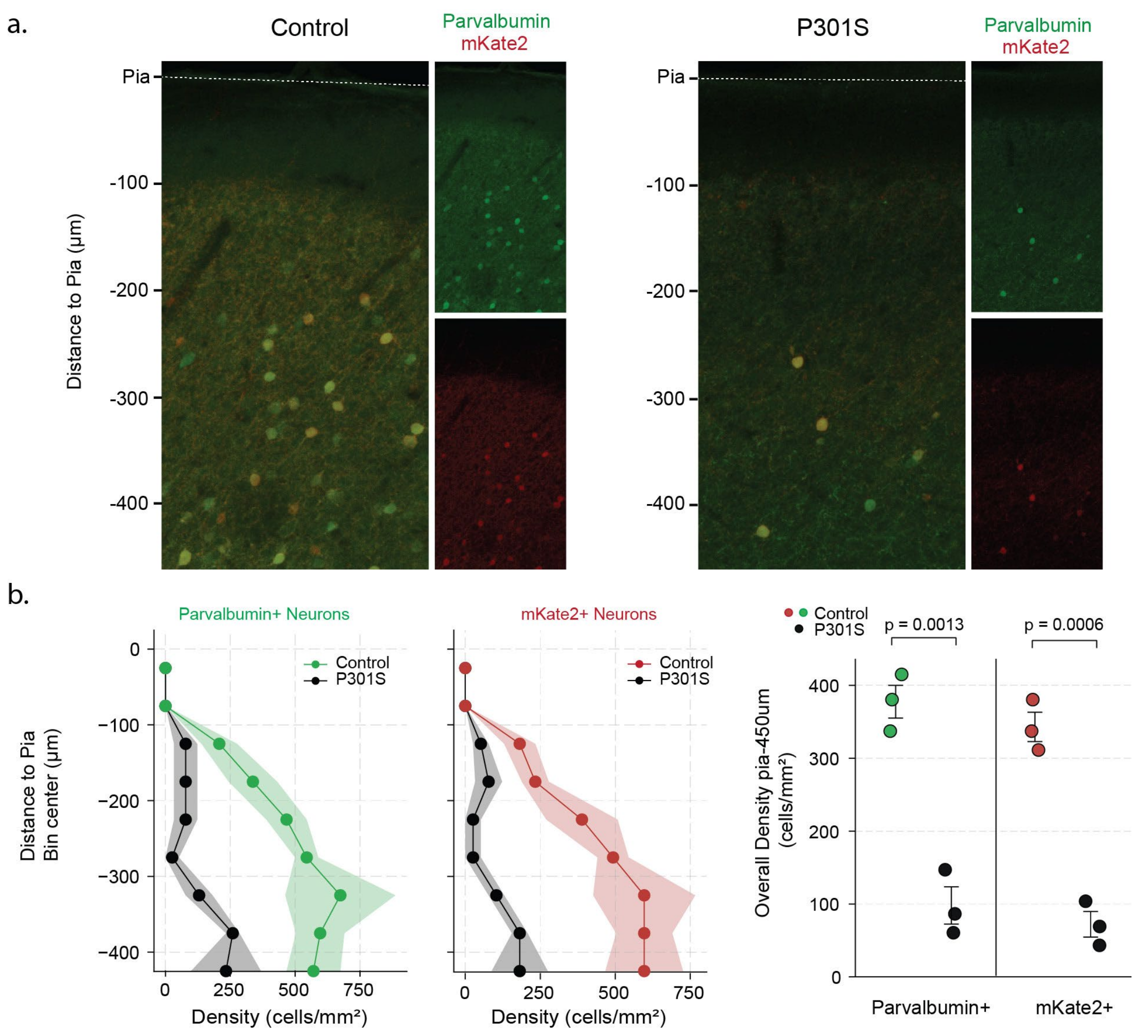


**Sup. Fig 3. (a)** Density of PV neurons in 6 months post injection mice was assessed at the center of the injection site via immunostaining against Parvalbumin (PV) and mKate2 protein. **(b)** Both the cell-type marker (PV), and the fluorescent label delivered via AAV (mKate2) were significantly reduced in the MAPT-P301S mice (N=3 mice per group, Welch's unpaired two-tailed t-test $p=0.0013$ and 0.0006 respectively).

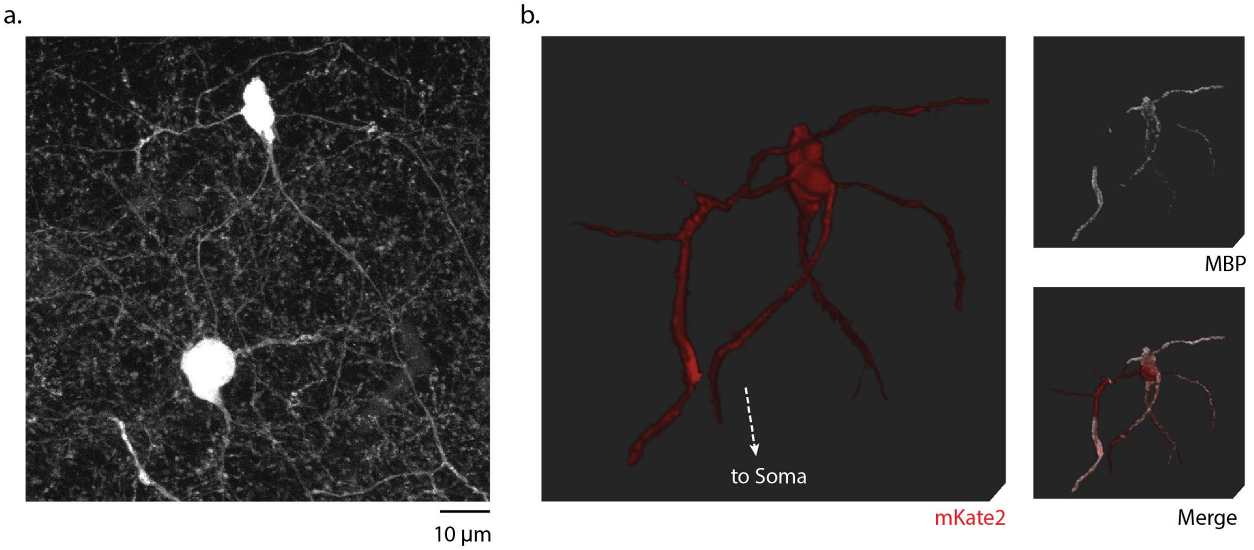


**Sup. Fig 4. (a)** An overview image showing a MAPT-P301S:mKate2 expressing Parvalbumin soma (at bottom of image), with an intact neurite projecting to a dystrophy (at top of image). Overview image shows the co-expressed mKate2 rendered in grayscale. **(b)** A separate high-resolution z-stack centered on the dystrophy presented in (a), visualized as a 3D rendering with mKate2 in red and MBP in white. Note the multiple emanating branches, and MBP positivity at the core of swelling, and along emanating branches.

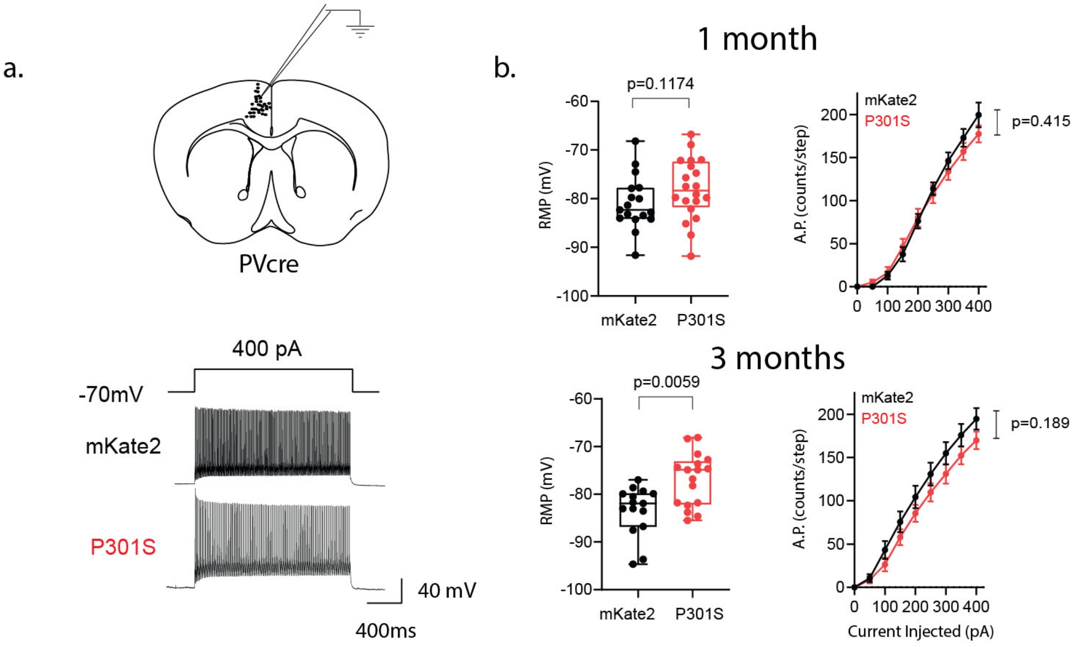


**Sup. Fig 5** *(***a)** PV neurons expressing control or tauopathy construct were recorded for electrophysiological characterization, showing example responses to current step injections. **(b)** Quantification of the resting membrane potential and elicited firing of PV neurons 1- and 3-months post injection (1M; P301S n=20 cells N=4 mice, control n=17 N=3. 3M; P301S n=17 N=3, control n=15, N=3). PV-P301S neurons displayed depolarized resting membrane potentials ($RMP_{1M}$; Mann-Whitney two-tailed p=0.1174; $RMP_{3M}$; Mann-Whitney two-tailed p=0.0059). Firing input–output relationships did not differ significantly between groups at either time point (1M: F = 2.96, neuron level permutation p = 0.415; 3M: F = 7.99, neuron level permutation p = 0.189).

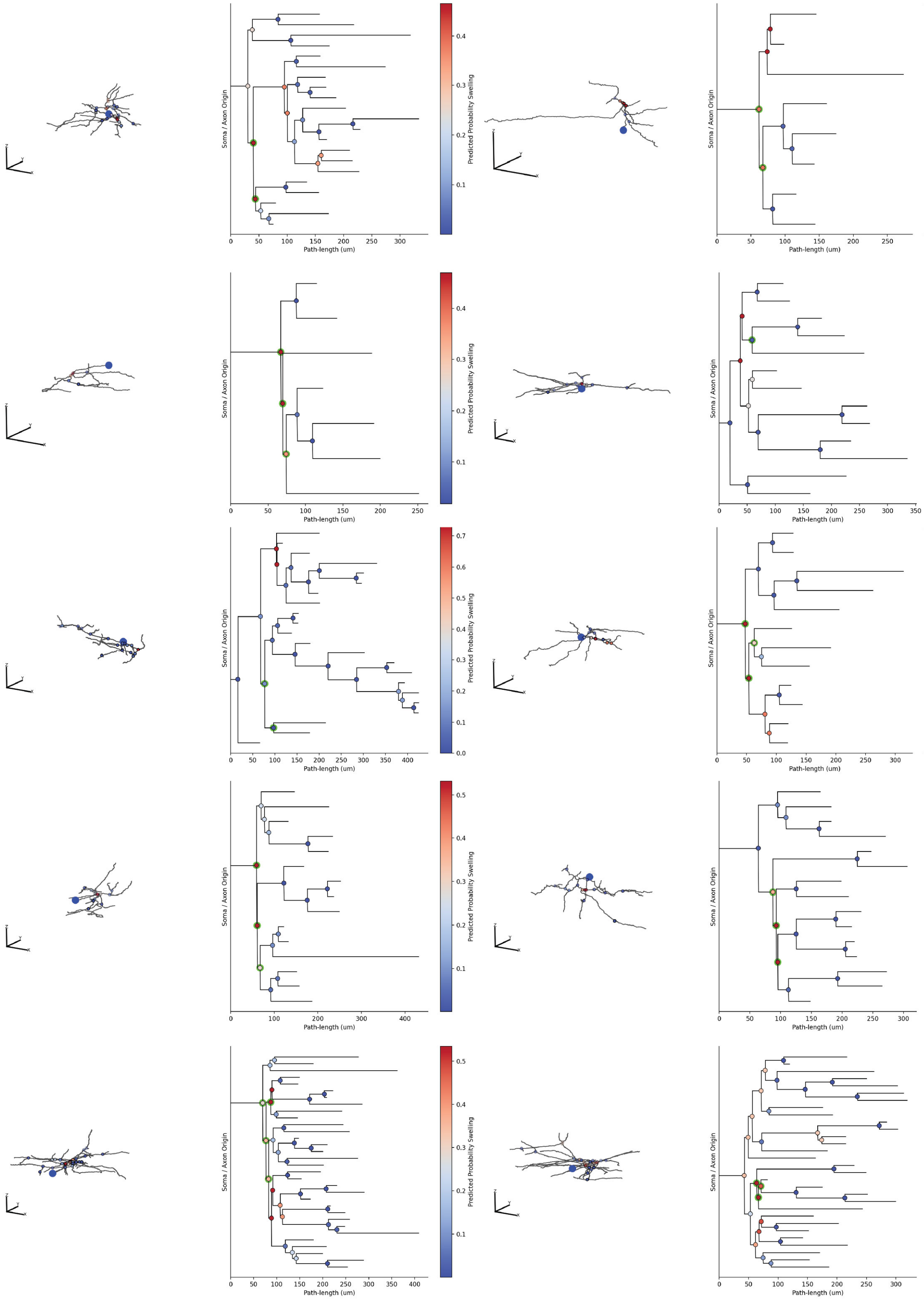


**Sup. Fig 6.** Generated axonograms of PV neurons expressing tauopathy construct. Left inserts show a 3D rendering of the reconstructed axon, large blue circle represents soma location. Right insert showing axonogram, with bifurcation colors corresponding to the predicted dystrophy probability according to the inter-bifurcation distance model generated by leave-one-neuron-out approach (color range scaled within each arbor for visualization, see main text, methods). Actual swelling locations are indicated by green highlights. 3D scalebar length 50 µm for all axes.

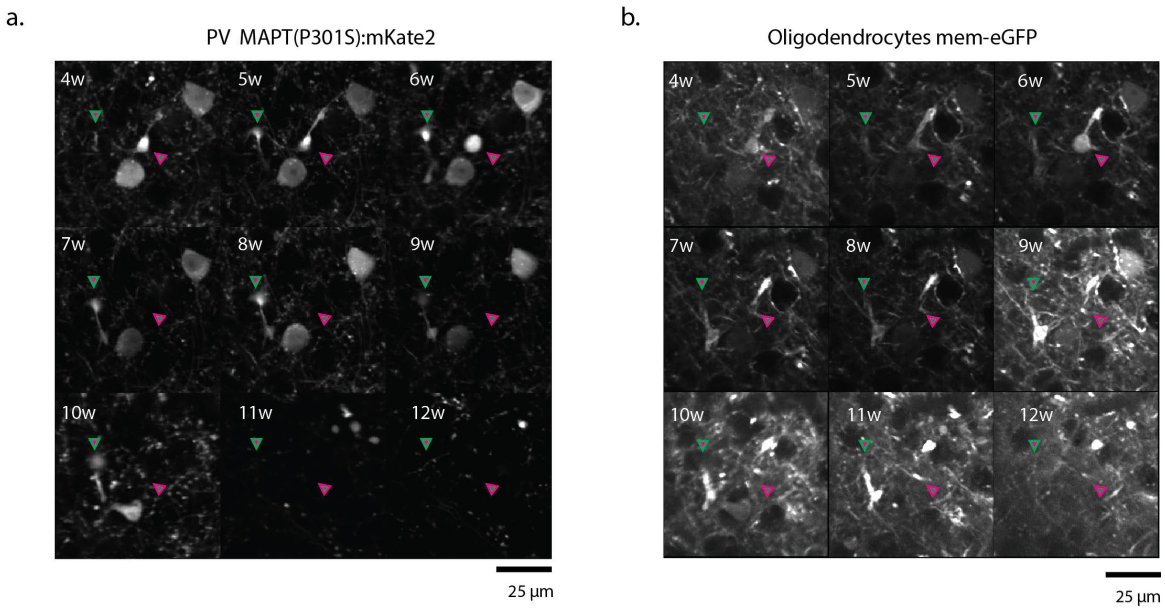


**Sup. Fig 7.** Time course of simultaneously imaged PV neuronal dystrophy and eGFP in oligodendrocyte membranes. Parvalbumin neurons expressed the P301S-mKate2 construct, and oligodendrocytes a membrane targeted eGFP (see methods). Magenta outlined arrows denote PV axonal swelling present at start of imaging, green outlined arrow denotes swelling emerging during time course acquisition.

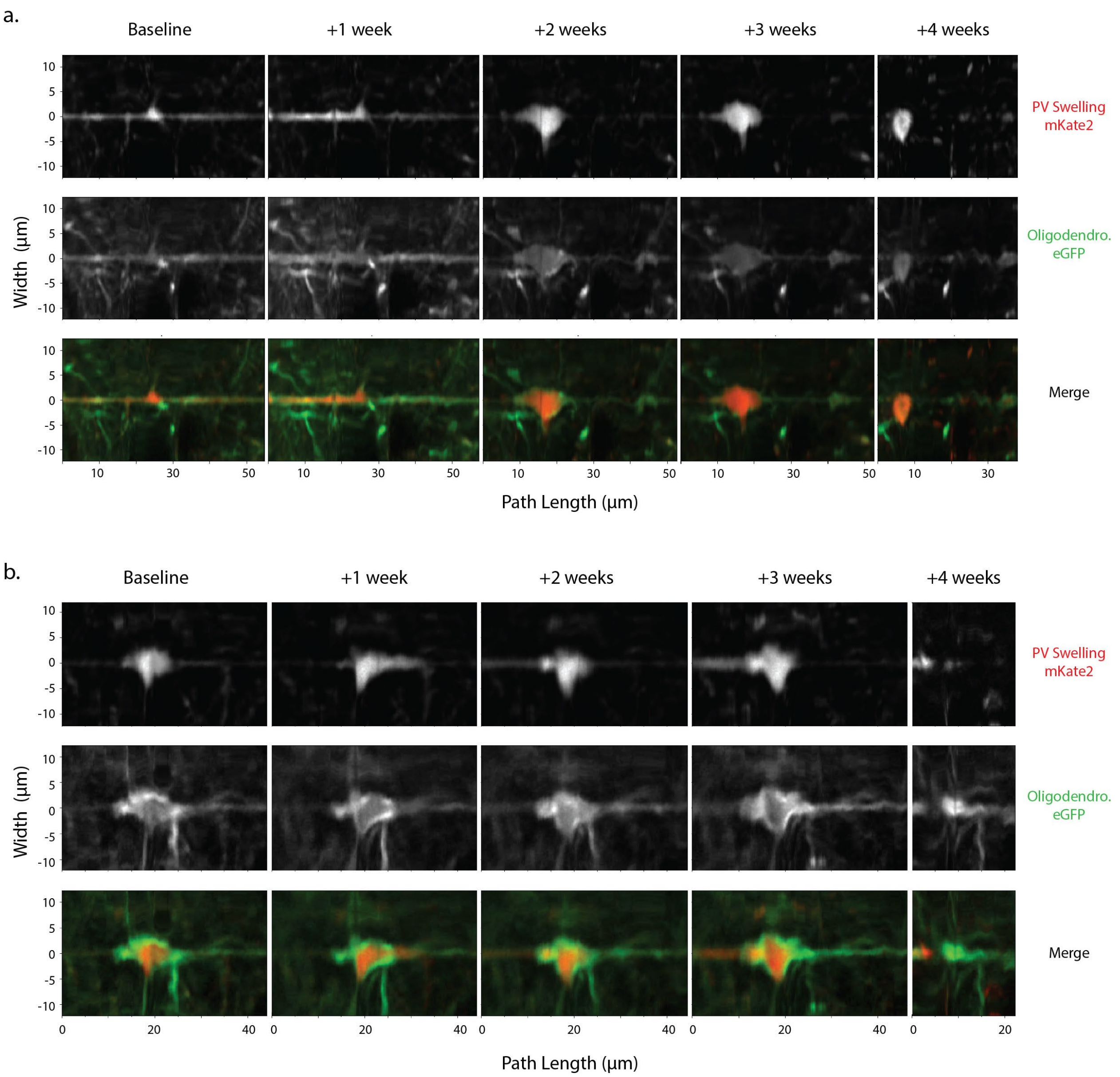


**Sup. Fig 8. (a,b)** Two example time-courses of progressive swelling of axon and Oligodendrocytic processes, images are digitally straightened along the axon path for visualization. The same location was recorded at one-week intervals. Parvalbumin neurons expressed the P301S-mKate2 construct, and oligodendrocytes a membrane targeted eGFP (see methods). The 2-color channel 3D volume was digitally straightened along a continuous portion of a PV axon path.

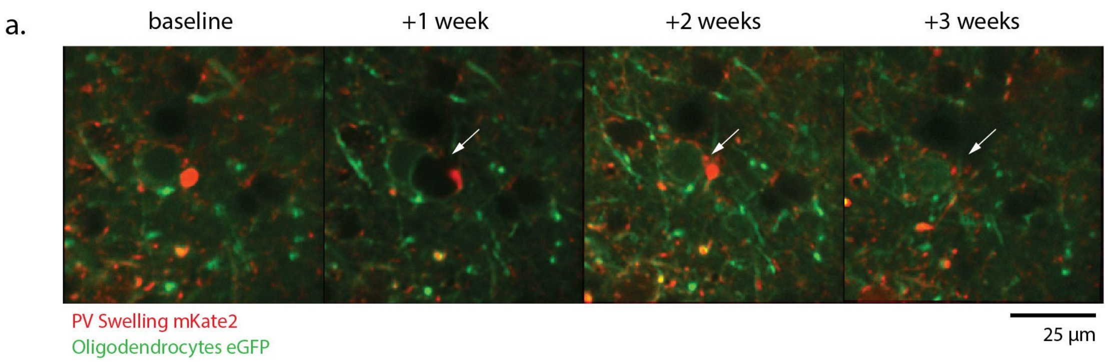


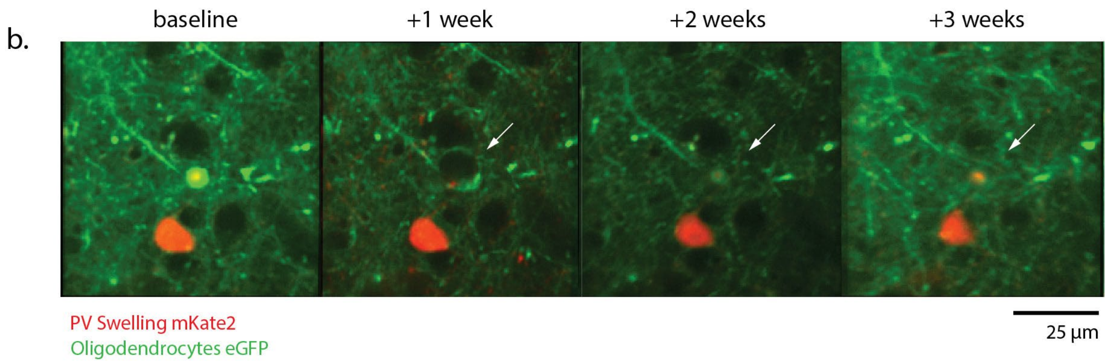


**Sup. Fig 9. (a,b)** Two example time-courses showing an acute expansion at +1 week relative to baseline timepoint. Note the recovery of oligodendrocyte morphology in subsequent timepoints. Note also the lack of correspondence to the PV+ axonal dystrophy morphology.

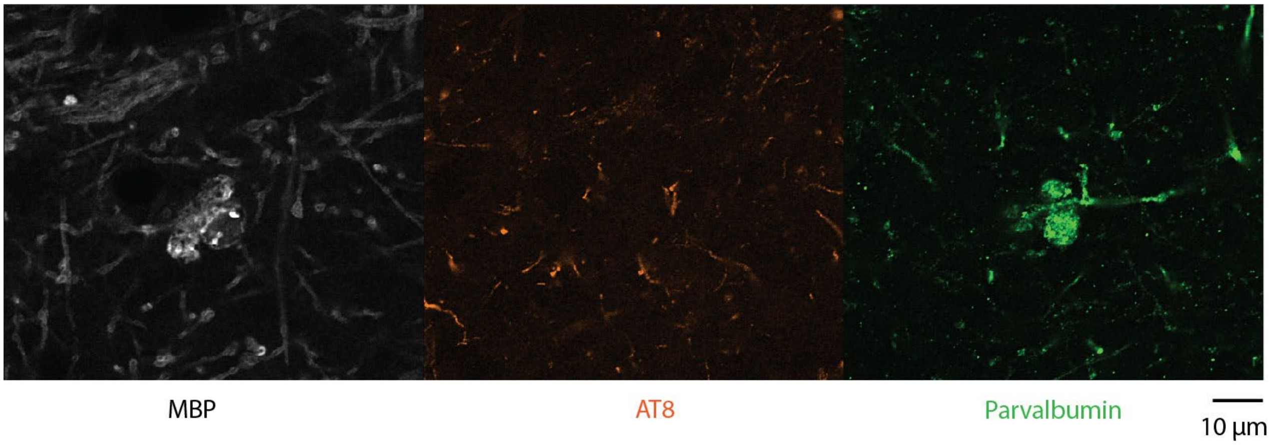


**Sup. Fig 10.** Single optical section image of a swelling at a PV axon bifurcation with severe accompanying myelin dystrophy. Image acquired in section from cortex of the superior frontal gyrus of a CBD patient. Immunostaining was performed against MBP (white), AT8 (orange), and Parvalbumin (green).

**Sup. Video 1**. 3D reconstruction centered on an axonal bifurcation associated dystrophy of a PV neuron expressing MAPT-P301S:mKate2. The video shows mKate2 (structural marker) and MBP (myelin). Note multiple emanating branches, and locally expanded myelin at the swelling.

**Vector Sequence (hTau-P301S:mKate2) for generation of AAV-EF1A-DIO-MAPT(P301S)-P2A-mKate2-WPRE**

CCTGCAGGCAGCTGCGCGCTCGCTCGCTCACTGAGGCCGCCCGGGCAAAGCCCGGGCGTCGGGCGACCTTTGGTC
GCCCGGCCTCAGTGAGCGAGCGAGCGCGCAGAGAGGGAGTGGCCAACTCCATCACTAGGGGTTCCTTCTAGACAA
CTTTGTATAGAAAAGTTGGGCTCCGGTGCCCGTCAGTGGGCAGAGCGCACATCGCCCACAGTCCCCGAGAAGTTG
GGGGGAGGGGTCGGCAATTGAACCGGTGCCTAGAGAAGGTGGCGCGGGGTAAACTGGGAAAGTGATGTCGTGTA
CTGGCTCCGCCTTTTTCCCGAGGGTGGGGGAGAACCGTATATAAGTGCAGTAGTCGCCGTGAACGTTCTTTTTCGC
AACGGGTTTGCCGCCAGAACACAGGTAAGTGCCGTGTGTGGTTCCCGCGGGCCTGGCCTCTTTACGGGTTATGGCC
CTTGCGTGCCTTGAATTACTTCCACCTGGCTGCAGTACGTGATTCTTGATCCCGAGCTTCGGGTTGGAAGTGGGTGG
GAGAGTTCGAGGCCTTGCGCTTAAGGAGCCCCTTCGCCTCGTGCTTGAGTTGAGGCCTGGCCTGGGCGCTGGGGCC
GCCGCGTGCGAATCTGGTGGCACCTTCGCGCCTGTCTCGCTGCTTTCGATAAGTCTCTAGCCATTTAAAATTTTTGA
TGACCTGCTGCGACGCTTTTTTTCTGGCAAGATAGTCTTGTAAATGCGGGCCAAGATCTGCACACTGGTATTTCGGT
TTTTGGGGCCGCGGGCGGCGACGGGGCCCGTGCGTCCCAGCGCACATGTTCGGCGAGGCGGGGCCTGCGAGCGCG
GCCACCGAGAATCGGACGGGGGTAGTCTCAAGCTGGCCGGCCTGCTCTGGTGCCTGGTCTCGCGCCGCCGTGTATC
GCCCCGCCCTGGGCGGCAAGGCTGGCCCGGTCGGCACCAGTTGCGTGAGCGGAAAGATGGCCGCTTCCCGGCCCT
GCTGCAGGGAGCTCAAAATGGAGGACGCGGCGCTCGGGAGAGCGGGCGGGTGAGTCACCCACACAAAGGAAAAG
GGCCTTTCCGTCCTCAGCCGTCGCTTCATGTGACTCCACGGAGTACCGGGCGCCGTCCAGGCACCTCGATTAGTTCT
CGAGCTTTTGGAGTACGTCGTCTTTAGGTTGGGGGGAGGGGTTTTATGCGATGGAGTTTCCCCACACTGAGTGGGT
GGAGACTGAAGTTAGGCCAGCTTGGCACTTGATGTAATTCTCCTTGGAATTTGCCCTTTTTGAGTTTGGATCTTGGT
TCATTCTCAAGCCTCAGACAGTGGTTCAAAGTTTTTTTCTTCCATTTCAGGTGTCGTGACAAGTTTGTACAAAAAAG
CAGGCTATAACTTCGTATAGGATACTTTATACGAAGTTATCCTCGAGTACCGGTCACGCGTACGCAAGCTTGATAA
CTTCGTATAGCATACATTATACGAAGTTATCCTAGGGCCACCTCACAAACCCTGCTTGGCCAGGGAGGCAGACACC
TCGTCAGCTAGCGTGGCGAGCTGGGGCGAGTCTACCATGTCGATGCTGCCGGTGGAGGAGACATTGCTGAGATGC
CGTGGAGACGTGTCCCCAGACACCACTGGCGACTTGTACACGATCTCCGCCCCGTGGTCTGTCTTGGCTTTGGCGT
TCTCGCGGAAGGTCAGCTTGTGGGTTTCAATCTTTTTATTTCCTCCGCCAGGGACGTGGGTGATATTGTCCAGGGAC
CCAATCTTCGACTGGACTCTGTCCTTGAAGTCAAGCTTCTCAGATTTTACTTCCACCTGGCCACCTCCTGGTTTATG
ATGGATGTTGCCTAATGAGCCACACTTGGAGGTCACCTTGCTCAGGTCAACTGGTTTGTAGACTATTTGCACACTG
CCGCCTCCGCTGACGTGTTTGATATTATCCTTTGAGCCACACTTGGACTGGACGTTGCTAAGATCCAGCTTCTTATT
AATTATCTGCACCTTCCCGCCTCCCGGCTGGTGCTTCAGGTTCTCAGTGGAGCCGATCTTGGACTTGACATTCTTCA
GGTCTGGCATGGGCACGGGGGCTGTCTGCAGGCGGCTCTTGGCGGAAGACGGCGACTTGGGTGGAGTACGGACCA
CTGCCACCTTCTTGGGCTCCCGGGTGGGTGGGGTTGGAAGGGACGGGGTGCGGGAGCGGCTGCCGGGAGTGCCTG
GGGAGCCGGGGCTGCTGTAGCCGCTGCGATCCCCTGATTTTGGAGGTTCACCAGAGCTGGGTGGTGTCTTTGGAGC
GGGCGGGGTTTTTGCTGGAATCCTGGTGGCGTTGGCCTGGCCCTTCTGGCCTGGAGGGGCTGCTCCCCGCGGTGTG
GCGATCTTCGTTTTACCATCAGCCCCCTTGGCTTTTTTGTCATCGCTTCCAGTCCCGTCTTTGCTTTTACTGACCATG
CGAGCTTGGGTCACGTGACCAGCAGCTTCGTCTTCCAGGCTGGGGGTGTCTCCAATGCCTGCTTCTTCAGCTGTGGT
TCCTTCTGGGATCTCCGTGTGGGGCTGCGCGGCAGCCTGCTTGCCGGGAGCTCCCTCATCCACTAAGGGTGCTGTC
ACATCTTCCGCTGTTGGAGTGCTCTTAGCATCAGAGGTTTCAGAGCCCGGTTCCTCAGATCCGTCCTCAGTGGGGGT
CTGCAGGGGAGATTCTTTCAGGCCAGCGTCCGTGTCACCCTCTTGGTCTTGGTGCATGGTGTAGCCCCCCTGATCTT
TCCTGTCCCCCAACCCGTACGTCCCAGCGTGATCTTCCATCACTTCGAACTCCTGGCGGGGCTCAGCCATAGGCCC
GGGGTTTCTTCAACATCTCCTGCTTGCTTTAACAGAGAGAAGTTCGTGGCTCCGCTTCCTCTGTGCCCCAGTTTGC
TAGGGAGGTCGCAGTATCTGGCCACAGCCACCTCGTGCTGCTCGACGTAGGTCTCTTTGTCGGCCTCCTTGATTCTT
TCCAGTCTTCTGTCCACATAGTAGACGCCGGGCATCTTGAGGTTCTTAGCGGGTTTCTTGGATCTGTATGTGGTCTT
CAAGTTGCAGATCAGGTGGCCCCCGCCCACGAGCTTCAGGGCCATGTCGGCTCTGCCTTCCAGGCCGCCGTCAGCG
GGGTACAGGGTCTCGGTGGAGGCCTCCCAGCCGAGTGTTTTCTTCTGCATCACAGGGCCGTTGGATGGGAAGTTCA
CCCCTCTGATCTTGACGTTGTAGATGAGGCAGCCGTCCTGGAGGCTGGTGTCCTGGGTAGCGGTCAGCACGCCCCC
GTCTTCGTATGTGGTGACTCTCTCCCATGTGAAGCCCTCGGGGAAGGACTGCTTAAAGAAGTCGGGGATGCCCTGG
GTGTGGTTGATGAAGGTTTTGCTGCCGTACATGAAGCTGGTAGCCAGGATGTCGAAGGCGAAGGGGAGAGGGCCG
CCCTCGACCGCCTTGATTCTCATGGTCTGGGTGCCCTCGTAGGGCTTGCCTTCGCCCTCGGATGTGCACTTGAAGTG
GTGGTTGTTCACGGTGCCCTCCATGTACAGCTTCATGTGCATGTTCTCCTTAATCAGCTCGCTCACCATGGTGGCTC
TAGAATAACTTCGTATAAAGTATCCTATACGAAGTTATAGGTACCTGAGCTCAACATCGATAGCACTAGTATAACT
TCGTATAATGTATGCTATACGAAGTTATACCCAGCTTTCTTGTACAAAGTGGGAATTCCGATAATCAACCTCTGGAT
TACAAAATTTGTGAAAGATTGACTGGTATTCTTAACTATGTTGCTCCTTTTACGCTATGTGGATACGCTGCTTTAAT
GCCTTTGTATCATGCTATTGCTTCCCGTATGGCTTTCATTTTCTCCTCCTTGTATAAATCCTGGTTGCTGTCTCTTTAT
GAGGAGTTGTGGCCCGTTGTCAGGCAACGTGGCGTGGTGTGCACTGTGTTTGCTGACGCAACCCCCACTGGTTGGG
GCATTGCCACCACCTGTCAGCTCCTTTCCGGGACTTTCGCTTTCCCCCTCCCTATTGCCACGGCGGAACTCATCGCC
GCCTGCCTTGCCCGCTGCTGGACAGGGGCTCGGCTGTTGGGCACTGACAATTCCGTGGTGTTGTCGGGGAAGCTGA
CGTCCTTTCCATGGCTGCTCGCCTGTGTTGCCACCTGGATTCTGCGCGGGACGTCCTTCTGCTACGTCCCTTCGGCC
CTCAATCCAGCGGACCTTCCTTCCCGCGGCCTGCTGCCGGCTCTGCGGCCTCTTCCGCGTCTTCGCCTTCGCCCTCA
GACGAGTCGGATCTCCCTTTGGGCCGCCTCCCCGCATCGGGAATTCCTAGAGCTCGCTGATCAGCCTCGACTGTGC
CTTCTAGTTGCCAGCCATCTGTTGTTTGCCCCTCCCCCGTGCCTTCCTTGACCCTGGAAGGTGCCACTCCCACTGTC

CTTTCCTAATAAAATGAGGAAATTGCATCGCATTGTCTGAGTAGGTGTCATTCTATTCTGGGGGGTGGGGTGGGGC
AGGACAGCAAGGGGGAGGATTGGGAAGAGAATAGCAGGCATGCTGGGGAGGGCCGCAGGAACCCCTAGTGATGG
AGTTGGCCACTCCCTCTCTGCGCGCTCGCTCGCTCACTGAGGCCGGGCGACCAAAGGTCGCCCGACGCCCGGGCTT
TGCCCGGGCGGCCTCAGTGAGCGAGCGAGCGCGCAGCTGCCTGCAGGGGCGCCTGATGCGGTATTTTCTCCTTACG
CATCTGTGCGGTATTTCACACCGCATACGTCAAAGCAACCATAGTACGCGCCCTGTAGCGGCGCATTAAGCGCGGC
GGGGGTGGTGGTTACGCGCAGCGTGACCGCTACACTTGCCAGCGCCTTAGCGCCCGCTCCTTTCGCTTTCTTCCCTT
CCTTTCTCGCCACGTTCGCCGGCTTTCCCCGTCAAGCTCTAAATCGGGGGCTCCCTTTAGGGTTCCGATTTAGTGCT
TTACGGCACCTCGACCCCAAAAAACTTGATTTGGGTGATGGTTCACGTAGTGGGCCATCGCCCTGATAGACGGTTT
TTCGCCCTTTGACGTTGGAGTCCACGTTCTTTAATAGTGGACTCTTGTTCCAAACTGGAACAACACTCAACTCTATC
TCGGGCTATTCTTTTGATTTATAAGGGATTTTGCCGATTTCGGTCTATTGGTTAAAAAATGAGCTGATTTAACAAAA
ATTTAACGCGAATTTTAACAAAATATTAACGTTTACAATTTTATGGTGCACTCTCAGTACAATCTGCTCTGATGCCG
CATAGTTAAGCCAGCCCCGACACCCGCCAACACCCGCTGACGCGCCCTGACGGGCTTGTCTGCTCCCGGCATCCGC
TTACAGACAAGCTGTGACCGTCTCCGGGAGCTGCATGTGTCAGAGGTTTTCACCGTCATCACCGAAACGCGCGAGA
CGAAAGGGCCTCGTGATACGCCTATTTTTATAGGTTAATGTCATGATAATAATGGTTTCTTAGACGTCAGGTGGCA
CTTTTCGGGGAAATGTGCGCGGAACCCCTATTTGTTTATTTTTCTAAATACATTCAAATATGTATCCGCTCATGAGA
CAATAACCCTGATAAATGCTTCAATAATATTGAAAAAGGAAGAGTATGAGTATTCAACATTTCCGTGTCGCCCTTA
TTCCCTTTTTTGCGGCATTTTGCCTTCCTGTTTTTGCTCACCCAGAAACGCTGGTGAAAGTAAAAGATGCTGAAGAT
CAGTTGGGTGCACGAGTGGGTTACATCGAACTGGATCTCAACAGCGGTAAGATCCTTGAGAGTTTTCGCCCCGAAG
AACGTTTTCCAATGATGAGCACTTTTAAAGTTCTGCTATGTGGCGCGGTATTATCCCGTATTGACGCCGGGCAAGA
GCAACTCGGTCGCCGCATACACTATTCTCAGAATGACTTGGTTGAGTACTCACCAGTCACAGAAAAGCATCTTACG
GATGGCATGACAGTAAGAGAATTATGCAGTGCTGCCATAACCATGAGTGATAACACTGCGGCCAACTTACTTCTGA
CAACGATCGGAGGACCGAAGGAGCTAACCGCTTTTTTGCACAACATGGGGGATCATGTAACTCGCCTTGATCGTTG
GGAACCGGAGCTGAATGAAGCCATACCAAACGACGAGCGTGACACCACGATGCCTGTAGCAATGGCAACAACGTT
GCGCAAACTATTAACTGGCGAACTACTTACTCTAGCTTCCCGGCAACAATTAATAGACTGGATGGAGGCGGATAA
AGTTGCAGGACCACTTCTGCGCTCGGCCCTTCCGGCTGGCTGGTTTATTGCTGATAAATCTGGAGCCGGTGAGCGT
GGAAGCCGCGGTATCATTGCAGCACTGGGGCCAGATGGTAAGCCCTCCCGTATCGTAGTTATCTACACGACGGGG
AGTCAGGCAACTATGGATGAACGAAATAGACAGATCGCTGAGATAGGTGCCTCACTGATTAAGCATTGGTAACTG
TCAGACCAAGTTTACTCATATATACTTTAGATTGATTTAAAACTTCATTTTTAATTTAAAAGGATCTAGGTGAAGAT
CCTTTTTGATAATCTCATGACCAAAATCCCTTAACGTGAGTTTTCGTTCCACTGAGCGTCAGACCCCGTAGAAAAG
ATCAAAGGATCTTCTTGAGATCCTTTTTTTCTGCGCGTAATCTGCTGCTTGCAAACAAAAAAACCACCGCTACCAGC
GGTGGTTTGTTTGCCGGATCAAGAGCTACCAACTCTTTTTCCGAAGGTAACTGGCTTCAGCAGAGCGCAGATACCA
AATACTGTTCTTCTAGTGTAGCCGTAGTTAGGCCACCACTTCAAGAACTCTGTAGCACCGCCTACATACCTCGCTCT
GCTAATCCTGTTACCAGTGGCTGCTGCCAGTGGCGATAAGTCGTGTCTTACCGGGTTGGACTCAAGACGATAGTTA
CCGGATAAGGCGCAGCGGTCGGGCTGAACGGGGGGTTCGTGCACACAGCCCAGCTTGGAGCGAACGACCTACACC
GAACTGAGATACCTACAGCGTGAGCTATGAGAAAGCGCCACGCTTCCCGAAGGGAGAAAGGCGGACAGGTATCC
GGTAAGCGGCAGGGTCGGAACAGGAGAGCGCACGAGGGAGCTTCCAGGGGGAAACGCCTGGTATCTTTATAGTCC
TGTCGGGTTTCGCCACCTCTGACTTGAGCGTCGATTTTTGTGATGCTCGTCAGGGGGGCGGAGCCTATGGAAAAAC
GCCAGCAACGCGGCCTTTTTACGGTTCCTGGCCTTTTGCTGGCCTTTTGCTCACATGT

**Vector Sequence (mKate2 control) for generation of AAV-EF1A-DIO-mKate2-WPRE**

CTGCGCGCTCGCTCGCTCACTGAGGCCGCCCGGGCAAAGCCCGGGCGTCGGGCGACCTTTGGTCGCCCGGCCTCAG
TGAGCGAGCGAGCGCGCAGAGAGGGAGTGGCCAACTCCATCACTAGGGGTTCCTTCTAGACAACTTTGTATAGAA
AAGTTGGGCTCCGGTGCCCGTCAGTGGGCAGAGCGCACATCGCCCACAGTCCCCGAGAAGTTGGGGGGAGGGGTC
GGCAATTGAACCGGTGCCTAGAGAAGGTGGCGCGGGGTAAACTGGGAAAGTGATGTCGTGTACTGGCTCCGCCTT
TTTCCCGAGGGTGGGGGAGAACCGTATATAAGTGCAGTAGTCGCCGTGAACGTTCTTTTTCGCAACGGGTTTGCCG
CCAGAACACAGGTAAGTGCCGTGTGTGGTTCCCGCGGGCCTGGCCTCTTTACGGGTTATGGCCCTTGCGTGCCTTG
AATTACTTCCACCTGGCTGCAGTACGTGATTCTTGATCCCGAGCTTCGGGTTGGAAGTGGGTGGGAGAGTTCGAGG
CCTTGCGCTTAAGGAGCCCCTTCGCCTCGTGCTTGAGTTGAGGCCTGGCCTGGGCGCTGGGGCCGCCGCGTGCGAA
TCTGGTGGCACCTTCGCGCCTGTCTCGCTGCTTTCGATAAGTCTCTAGCCATTTAAAATTTTTGATGACCTGCTGCG
ACGCTTTTTTTCTGGCAAGATAGTCTTGTAAATGCGGGCCAAGATCTGCACACTGGTATTTCGGTTTTTGGGGCCGC
GGGCGGCGACGGGGCCCGTGCGTCCCAGCGCACATGTTCGGCGAGGCGGGGCCTGCGAGCGCGGCCACCGAGAA
TCGGACGGGGGTAGTCTCAAGCTGGCCGGCCTGCTCTGGTGCCTGGTCTCGCGCCGCCGTGTATCGCCCCGCCCTG
GGCGGCAAGGCTGGCCCGGTCGGCACCAGTTGCGTGAGCGGAAAGATGGCCGCTTCCCGGCCCTGCTGCAGGGAG
CTCAAAATGGAGGACGCGGCGCTCGGGAGAGCGGGCGGGTGAGTCACCCACACAAAGGAAAAGGGCCTTTCCGT
CCTCAGCCGTCGCTTCATGTGACTCCACGGAGTACCGGGCGCCGTCCAGGCACCTCGATTAGTTCTCGAGCTTTTG
GAGTACGTCGTCTTTAGGTTGGGGGGAGGGGTTTTATGCGATGGAGTTTCCCCACACTGAGTGGGTGGAGACTGAA
GTTAGGCCAGCTTGGCACTTGATGTAATTCTCCTTGGAATTTGCCCTTTTTGAGTTTGGATCTTGGTTCATTCTCAAG
CCTCAGACAGTGGTTCAAAGTTTTTTTCTTCCATTTCAGGTGTCGTGACAAGTTTGTACAAAAAAGCAGGCTATAAC
TTCGTATAGGATACTTTATACGAAGTTATCCTCGAGTACCGGTCACGCGTACGCAAGCTTGATAACTTCGTATAGC
ATACATTATACGAAGTTATCCTAGGGCCACCTCATCTGTGCCCCAGTTTGCTAGGGAGGTCGCAGTATCTGGCCAC

AGCCACCTCGTGCTGCTCGACGTAGGTCTCTTTGTCGGCCTCCTTGATTCTTTCCAGTCTTCTGTCCACATAGTAGA
CGCCGGGCATCTTGAGGTTCTTAGCGGGTTTCTTGGATCTGTATGTGGTCTTCAAGTTGCAGATCAGGTGGCCCCCG
CCCACGAGCTTCAGGGCCATGTCGGCTCTGCCTTCCAGGCCGCCGTCAGCGGGGTACAGGGTCTCGGTGGAGGCCT
CCCAGCCGAGTGTTTTCTTCTGCATCACAGGGCCGTTGGATGGGAAGTTCACCCCTCTGATCTTGACGTTGTAGATG
AGGCAGCCGTCCTGGAGGCTGGTGTCCTGGGTAGCGGTCAGCACGCCCCCGTCTTCGTATGTGGTGACTCTCTCCC
ATGTGAAGCCCTCGGGGAAGGACTGCTTAAAGAAGTCGGGGATGCCCTGGGTGTGGTTGATGAAGGTTTTGCTGC
CGTACATGAAGCTGGTAGCCAGGATGTCGAAGGCGAAGGGGAGAGGGCCGCCCTCGACCGCCTTGATTCTCATGG
TCTGGGTGCCCTCGTAGGGCTTGCCTTCGCCCTCGGATGTGCACTTGAAGTGGTGGTTGTTCACGGTGCCCTCCATG
TACAGCTTCATGTGCATGTTCTCCTTAATCAGCTCGCTCACCATGGTGGCTCTAGAATAACTTCGTATAAAGTATCC
TATACGAAGTTATAGGTACCTGAGCTCAACATCGATAGCACTAGTATAACTTCGTATAATGTATGCTATACGAAGT
TATACCCAGCTTTCTTGTACAAAGTGGGAATTCCGATAATCAACCTCTGGATTACAAAATTTGTGAAAGATTGACT
GGTATTCTTAACTATGTTGCTCCTTTTACGCTATGTGGATACGCTGCTTTAATGCCTTTGTATCATGCTATTGCTTCC
CGTATGGCTTTCATTTTCTCCTCCTTGTATAAATCCTGGTTGCTGTCTCTTTATGAGGAGTTGTGGCCCGTTGTCAGG
CAACGTGGCGTGGTGTGCACTGTGTTTGCTGACGCAACCCCCACTGGTTGGGGCATTGCCACCACCTGTCAGCTCC
TTTCCGGGACTTTCGCTTTCCCCCTCCCTATTGCCACGGCGGAACTCATCGCCGCCTGCCTTGCCCGCTGCTGGACA
GGGGCTCGGCTGTTGGGCACTGACAATTCCGTGGTGTTGTCGGGGAAGCTGACGTCCTTTCCATGGCTGCTCGCCT
GTGTTGCCACCTGGATTCTGCGCGGGACGTCCTTCTGCTACGTCCCTTCGGCCCTCAATCCAGCGGACCTTCCTTCC
CGCGGCCTGCTGCCGGCTCTGCGGCCTCTTCCGCGTCTTCGCCTTCGCCCTCAGACGAGTCGGATCTCCCTTTGGGC
CGCCTCCCCGCATCGGGAATTCCTAGAGCTCGCTGATCAGCCTCGACTGTGCCTTCTAGTTGCCAGCCATCTGTTGT
TTGCCCCTCCCCCGTGCCTTCCTTGACCCTGGAAGGTGCCACTCCCACTGTCCTTTCCTAATAAAATGAGGAAATTG
CATCGCATTGTCTGAGTAGGTGTCATTCTATTCTGGGGGGTGGGGTGGGGCAGGACAGCAAGGGGGAGGATTGGG
AAGAGAATAGCAGGCATGCTGGGGAGGGCCGCAGGAACCCCTAGTGATGGAGTTGGCCACTCCCTCTCTGCGCGC
TCGCTCGCTCACTGAGGCCGGGCGACCAAAGGTCGCCCGACGCCCGGGCTTTGCCCGGGCGGCCTCAGTGAGCGA
GCGAGCGCGCAGCTGCCTGCAGGGGCGCCTGATGCGGTATTTTCTCCTTACGCATCTGTGCGGTATTTCACACCGC
ATACGTCAAAGCAACCATAGTACGCGCCCTGTAGCGGCGCATTAAGCGCGGCGGGGGTGGTGGTTACGCGCAGCG
TGACCGCTACACTTGCCAGCGCCTTAGCGCCCGCTCCTTTCGCTTTCTTCCCTTCCTTTCTCGCCACGTTCGCCGGCT
TTCCCCGTCAAGCTCTAAATCGGGGGCTCCCTTTAGGGTTCCGATTTAGTGCTTTACGGCACCTCGACCCCAAAAA
ACTTGATTTGGGTGATGGTTCACGTAGTGGGCCATCGCCCTGATAGACGGTTTTTCGCCCTTTGACGTTGGAGTCCA
CGTTCTTTAATAGTGGACTCTTGTTCCAAACTGGAACAACACTCAACTCTATCTCGGGCTATTCTTTTGATTTATAA
GGGATTTTGCCGATTTCGGTCTATTGGTTAAAAAATGAGCTGATTTAACAAAAATTTAACGCGAATTTTAACAAAA
TATTAACGTTTACAATTTTATGGTGCACTCTCAGTACAATCTGCTCTGATGCCGCATAGTTAAGCCAGCCCCGACAC
CCGCCAACACCCGCTGACGCGCCCTGACGGGCTTGTCTGCTCCCGGCATCCGCTTACAGACAAGCTGTGACCGTCT
CCGGGAGCTGCATGTGTCAGAGGTTTTCACCGTCATCACCGAAACGCGCGAGACGAAAGGGCCTCGTGATACGCC
TATTTTTATAGGTTAATGTCATGATAATAATGGTTTCTTAGACGTCAGGTGGCACTTTTCGGGGAAATGTGCGCGGA
ACCCCTATTTGTTTATTTTTCTAAATACATTCAAATATGTATCCGCTCATGAGACAATAACCCTGATAAATGCTTCA
ATAATATTGAAAAAGGAAGAGTATGAGTATTCAACATTTCCGTGTCGCCCTTATTCCCTTTTTTGCGGCATTTTGCC
TTCCTGTTTTTGCTCACCCAGAAACGCTGGTGAAAGTAAAAGATGCTGAAGATCAGTTGGGTGCACGAGTGGGTTA
CATCGAACTGGATCTCAACAGCGGTAAGATCCTTGAGAGTTTTCGCCCCGAAGAACGTTTTCCAATGATGAGCACT
TTTAAAGTTCTGCTATGTGGCGCGGTATTATCCCGTATTGACGCCGGGCAAGAGCAACTCGGTCGCCGCATACACT
ATTCTCAGAATGACTTGGTTGAGTACTCACCAGTCACAGAAAAGCATCTTACGGATGGCATGACAGTAAGAGAATT
ATGCAGTGCTGCCATAACCATGAGTGATAACACTGCGGCCAACTTACTTCTGACAACGATCGGAGGACCGAAGGA
GCTAACCGCTTTTTTGCACAACATGGGGGATCATGTAACTCGCCTTGATCGTTGGGAACCGGAGCTGAATGAAGCC
ATACCAAACGACGAGCGTGACACCACGATGCCTGTAGCAATGGCAACAACGTTGCGCAAACTATTAACTGGCGAA
CTACTTACTCTAGCTTCCCGGCAACAATTAATAGACTGGATGGAGGCGGATAAAGTTGCAGGACCACTTCTGCGCT
CGGCCCTTCCGGCTGGCTGGTTTATTGCTGATAAATCTGGAGCCGGTGAGCGTGGAAGCCGCGGTATCATTGCAGC
ACTGGGGCCAGATGGTAAGCCCTCCCGTATCGTAGTTATCTACACGACGGGGAGTCAGGCAACTATGGATGAACG
AAATAGACAGATCGCTGAGATAGGTGCCTCACTGATTAAGCATTGGTAACTGTCAGACCAAGTTTACTCATATATA
CTTTAGATTGATTTAAAACTTCATTTTTAATTTAAAAGGATCTAGGTGAAGATCCTTTTTGATAATCTCATGACCAA
AATCCCTTAACGTGAGTTTTCGTTCCACTGAGCGTCAGACCCCGTAGAAAAGATCAAAGGATCTTCTTGAGATCCT
TTTTTTCTGCGCGTAATCTGCTGCTTGCAAACAAAAAAACCACCGCTACCAGCGGTGGTTTGTTTGCCGGATCAAG
AGCTACCAACTCTTTTTCCGAAGGTAACTGGCTTCAGCAGAGCGCAGATACCAAATACTGTTCTTCTAGTGTAGCC
GTAGTTAGGCCACCACTTCAAGAACTCTGTAGCACCGCCTACATACCTCGCTCTGCTAATCCTGTTACCAGTGGCT
GCTGCCAGTGGCGATAAGTCGTGTCTTACCGGGTTGGACTCAAGACGATAGTTACCGGATAAGGCGCAGCGGTCG
GGCTGAACGGGGGGTTCGTGCACACAGCCCAGCTTGGAGCGAACGACCTACACCGAACTGAGATACCTACAGCGT
GAGCTATGAGAAAGCGCCACGCTTCCCGAAGGGAGAAAGGCGGACAGGTATCCGGTAAGCGGCAGGGTCGGAAC
AGGAGAGCGCACGAGGGAGCTTCCAGGGGGAAACGCCTGGTATCTTTATAGTCCTGTCGGGTTTCGCCACCTCTGA
CTTGAGCGTCGATTTTTGTGATGCTCGTCAGGGGGGCGGAGCCTATGGAAAAACGCCAGCAACGCGGCCTTTTTAC
GGTTCCTGGCCTTTTGCTGGCCTTTTGCTCACATGTCCTGCAGGCAG
